# Physiomimetic culture bias durotaxis toward soft environments

**DOI:** 10.64898/2026.03.24.713716

**Authors:** Marina Moro-López, Roberto Alonso-Matilla, Sergi Olivé, Manuel Gómez-González, Paolo Provenzano, Ramon Farré, Jorge Otero, David Odde, Raimon Sunyer

## Abstract

Directed cell migration underlies many biological phenomena, from embryonic development to tumor metastasis and organ fibrosis. Most cells typically migrate toward stiffer regions of their extracellular matrix –a behavior known as positive durotaxis. Here we show that culture on rigid plastic reinforces this response, whereas preconditioning in soft 3D physiomimetic environments reprograms migration towards softer environments, a phenomenon known as negative durotaxis. Fetal rat lung fibroblasts preconditioned in 3D physiomimetic hydrogels exhibited negative durotaxis and accumulated near ∼5 kPa, corresponding to the physiological stiffness of the lung. In contrast, genetically identical cells maintained on conventional 2D plastic substrates migrated up stiffness gradients, toward stiffer regions. Although both populations displayed a biphasic force-stiffness relationship, they differed in force magnitude and cytoskeletal organization. Molecular-clutch modeling revealed that durotaxis reversal emerges from two distinct mechanical regimes: a mechanosensitive, high-motor-clutch state that stabilizes adhesions on stiff substrates and drives positive durotaxis, and a low-motor, weak-adhesion state in which clutch slippage on the stiff side causes negative durotaxis. Our results show that durotaxis direction is not an intrinsic cellular property. Rather, it emerges from the interplay between motor activity and adhesion dynamics and can be tuned by culture conditions.

## Introduction

Directed cell migration is a fundamental biological process that shapes embryonic development^1^, sustains tissue homeostasis, and drives pathological events such as tumor invasion and organ fibrosis^2^. Directed migration is commonly attributed to chemotaxis, in which cells follow spatial gradients of chemical factors dissolved in the media. However, cells are also able to direct their migration by following spatial gradients in the mechanical properties of their extracellular matrix (ECM) –a phenomenon called durotaxis^3–5^. Durotaxis has been predominantly associated with migration toward stiffer regions, a process termed positive durotaxis^3,6–10^. However, this canonical view has been recently challenged. Some cell types exhibit reversed migratory responses, moving from stiff to soft regions and accumulating at intermediate rigidities^11^ –a process referred to as negative durotaxis. These observations raise the question of which cellular and mechanical factors govern the direction of durotaxis.

In this work, we combine experiments and theory to test whether durotaxis direction is shaped by culture conditions. From a physiological standpoint, it is unclear why most cells persistently migrate toward ever-stiffer regions via positive durotaxis. Instead, a more plausible scenario is that cells preferentially migrate toward the stiffness of their native tissue. Previous studies have shown that cell exposure to distinct mechanical environments induces persistent changes in cytoskeleton architecture^12,13^, transcriptional regulation^14,13,15^, invasiveness^13^, chromatin accessibility^16,17,15^, and mechanosensitivity^12,18^. We therefore asked whether prolonged culture on rigid, non-physiological plastic rewires migration to move up stiffness gradients, and whether physiomimetic 3D culture reverses that bias. We compared fetal rat fibroblasts preconditioned either on conventional plastic or in physiomimetic 3D lung-derived hydrogels. Although both populations exhibited a biphasic force-stiffness relationship, only physiomimetic-preconditioned cells displayed negative durotaxis, whereas plastic-cultured cells exhibited positive durotaxis. A motor-clutch-based Cell Migration Simulator^19^ explained this durotaxis reversal through stiffness-dependent motor activity and moderate adhesion reinforcement for plastic preconditioned cells. These results show that culturing conditions deeply reconfigure the cytoskeleton and motor machinery, biasing durotaxis toward soft or stiff environments.

## Results

To study how the culture environment modulates cell mechanoresponse and durotaxis, we used RFL-6 cells, a non-transformed fibroblast line derived from fetal lung tissue of Sprague-Dawley rats. These cells retain defining features of stromal fibroblasts, including mesenchymal morphology and sensitivity to ECM cues. Cells were cultured for four days either on standard tissue-culture plastic or within physiomimetic hydrogels derived from decellularized porcine lungs (Fig. 1a). Hydrogels were prepared by cryogenic milling, enzymatic digestion, and pH neutralization following established protocols^20^ (see Methods and Fig. 1a, inset). This approach provides a 3D scaffold that recapitulates biochemical and mechanical signals from the native environment of the lung. After preconditioning, cells from both conditions were transferred to collagen I-coated polyacrylamide (PAA) gels of defined stiffness (0.5 to 30kPa, see Methods, Supplementary Text I and Extended Data Fig. 1). On these substrates, we quantified cell spreading area, actomyosin organization, focal adhesion (FA) organization, migratory behavior, and traction forces (Fig. 1a).

**Figure 1.**
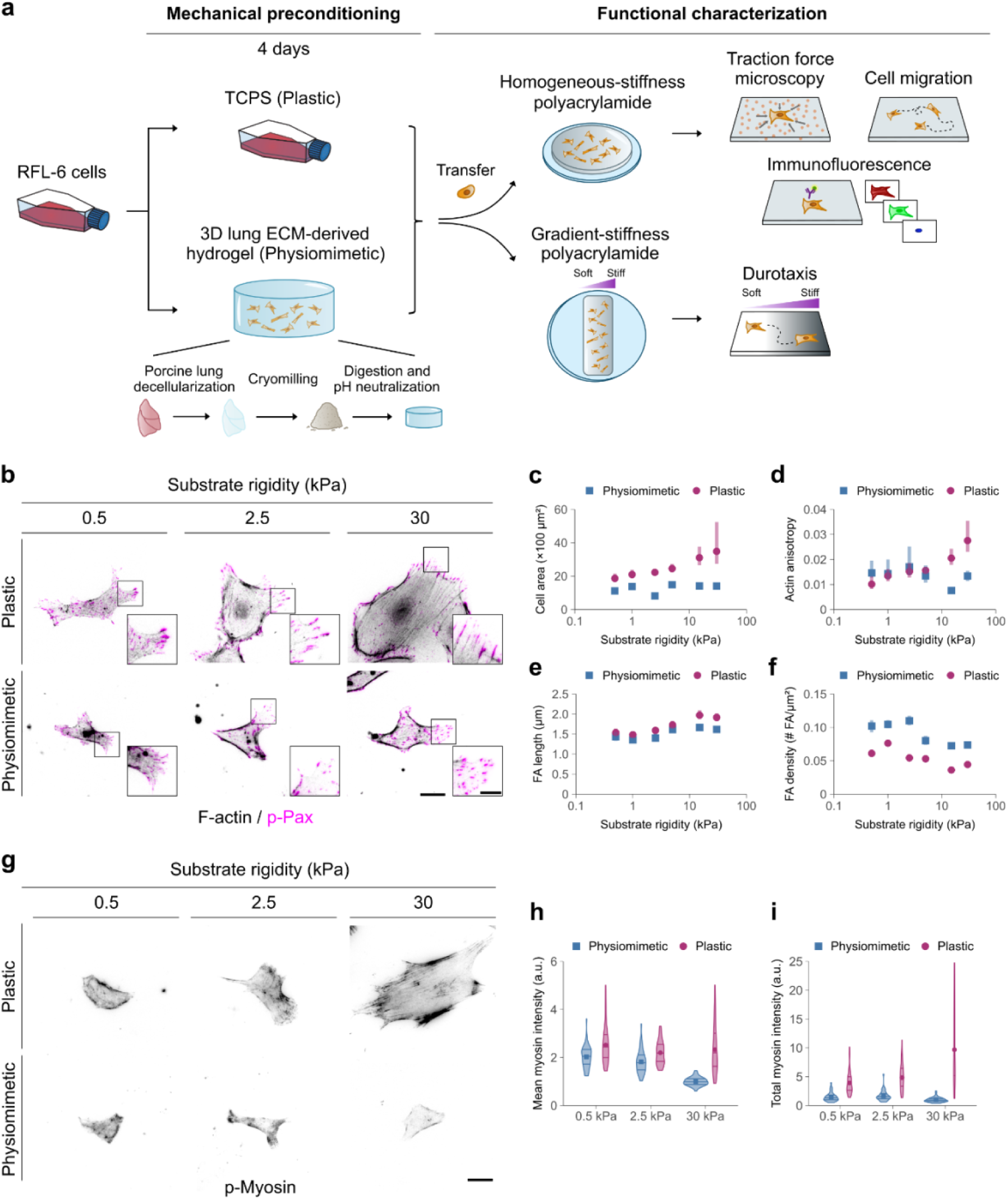
Plastic and physiomimetic preconditioning establish distinct cytoskeletal architecture. **a,** Experimental design. RFL-6 fibroblasts were cultured for 4 days on plastic or in physiomimetic hydrogels. After preconditioning, cells were replated on PAA gels (0.5–30 kPa) to quantify mechanoresponses. Inset, workflow of hydrogel preparation by cryogenic milling, enzymatic digestion, and pH neutralization. **b,** Representative images of actin filaments (phalloidin) and focal adhesions (FAs, p-Paxillin) in plastic- and physiomimetic-preconditioned cells on 0.5, 2.5 and 30 kPa substrates (scale bar, 25 µm). Inset, magnified view highlighting individual FAs (scale bar, 10 µm). **c,** Cell spreading area across substrate stiffness. Plastic-preconditioned cells displayed rigidity-dependent growth, whereas physiomimetic-preconditioned cells were largely insensitive to stiffness. **d,** Quantification of actin fiber alignment. Plastic-preconditioned cells exhibited a marked increase in alignment on stiff matrices, whereas physiomimetic-preconditioned cells did not. **e,** FA length as a function of substrate stiffness. Plastic-preconditioned cells consistently exhibited larger FAs and a steeper stiffness-dependent increase. **f,** FA density across substrate stiffness. Both groups showed a significant reduction in FA density at higher stiffness. For panels **c–f**, statistical comparisons were performed using a two-way permutation ANOVA. Both preconditioning and stiffness showed significant main effects (*p* < 0.0001 for all measures except FA length, where preconditioning *p* = 0.0459 and stiffness *p* = 0.0002). The interaction term was significant in all the panels, indicating that the influence of stiffness on cell spreading area **(c)**, actin alignment **(d)**, FA length **(e)** and FA density **(f)** differed between preconditioned groups. Data represent mean ± 95% confidence intervals, estimated by bootstrap resampling of 45 cells per stiffness and condition from three independent experiments. **g,** Representative images of phosphorylated myosin (p-Myosin) in plastic- and physiomimetic-preconditioned cells cultured on 0.5, 2.5 and 30 kPa substrates (scale bar, 25 µm). **h,** Mean p-Myosin normalized to the physiomimetic-30 kPa mean intensity. Cells preconditioned on plastic exhibited higher p-Myosin levels. **i,** Total p-Myosin intensity normalized to the physiomimetic-30 kPa total intensity. Plastic-preconditioned cells displayed a significant stiffness-dependent rise in p-Myosin, whereas physiomimetic-preconditioned cells showed a weaker response. For panels **(h)** and **(i)** a permutation-based ANOVA confirmed significant effects of condition, stiffness, and their interaction (*p* < 0.0001 for all terms). Data represent n = 45 cells from 3 independent experiments.

### Preconditioning shapes cytoskeletal architecture

We first examined how preconditioning on standard plastic or physiomimetic hydrogels modulated subsequent mechanoresponses on defined stiffness PAA gels. To isolate cell-matrix from cell-cell interactions, we analyzed single cells plated at low density. We imaged cells 4 h after attachment to minimize accumulation of newly deposited ECM. Actin filaments and FAs were visualized by phalloidin and phosphorylated paxillin (p-Pax) staining (see Methods). Plastic-preconditioned cells showed a pronounced response to increasing substrate stiffness, characterized by enhanced cell spreading (Fig. 1b,c), the assembly of thick, aligned stress fibers (Fig. 1b,d), and the elongation of FAs predominantly localized at the cell periphery (Fig. 1b,e and Extended Data Fig. 2). Surprisingly, although FAs elongated with increasing substrate stiffness, their density decreased by ∼50%, indicating that adhesions matured under tension but failed to scale with cell spreading (Fig. 1f). Additionally, phosphorylated myosin II (p-Myosin) levels increased with stiffness (Fig. 1g-i), suggesting that individual adhesions transmit large forces at high stiffness, but through fewer contact sites.

Physiomimetic-preconditioned cells, by contrast, were less sensitive to stiffness: spreading area remained constant across substrates (Fig. 1b,c), and the actin cytoskeleton was thin, poorly aligned, and disorganized (Fig. 1b,d). FAs elongation also increased with stiffness, albeit less markedly than in plastic-preconditioned cells (Fig. 1b,e), and peripheral accumulation was modest (Extended Data Fig. 2). The FA density decreased by 34% with increasing stiffness, suggesting a reduced capacity to maintain adhesion number at higher rigidities (Fig. 1f). p-Myosin levels were significantly lower than in plastic-preconditioned cells and mildly decreased with stiffness (Fig. 1g-i). Together, these results show that preconditioning on plastic or physiomimetic matrices establishes distinct cytoskeletal and adhesion architectures, leading to divergent stiffness-dependent regulation of spreading, force transmission, and actomyosin organization.

### Preconditioning modulates traction force

Given that cytoskeletal remodeling is closely linked to actomyosin contractility, we next assessed how these alterations translated into cell-substrate force generation (Fig. 2a). Traction forces were quantified across substrates of varying rigidity using traction force microscopy (see Methods). Two key features emerged. First, consistent with p-Myosin quantification (Fig. 1g-i), plastic-preconditioned cells exerted systematically higher traction forces –by 40–70% across the entire stiffness range– than physiomimetic-preconditioned cells (permutation ANOVA, *p* = 0.0001). Second, unlike the monotonic increase with stiffness described for many cell types^21–26^, both populations displayed biphasic, peak-shaped force-stiffness relationships. A quadratic permutation ANOVA confirmed a significant nonlinear dependence on stiffness (*p* = 0.0027 for the quadratic term), indicating that both conditions exert maximal force at intermediate stiffness. Total traction force for plastic-preconditioned cells peaked at ∼2.5 kPa and declined thereafter, whereas traction force for physiomimetic-preconditioned cells increased up to ∼2.5–5 kPa before dropping at higher stiffness (Fig. 2b,c). Together, these findings suggest that both cell populations are mechanically adapted to exert maximal force at an intermediate stiffness yet differ in their absolute force regime because of preconditioning.

**Figure 2.**
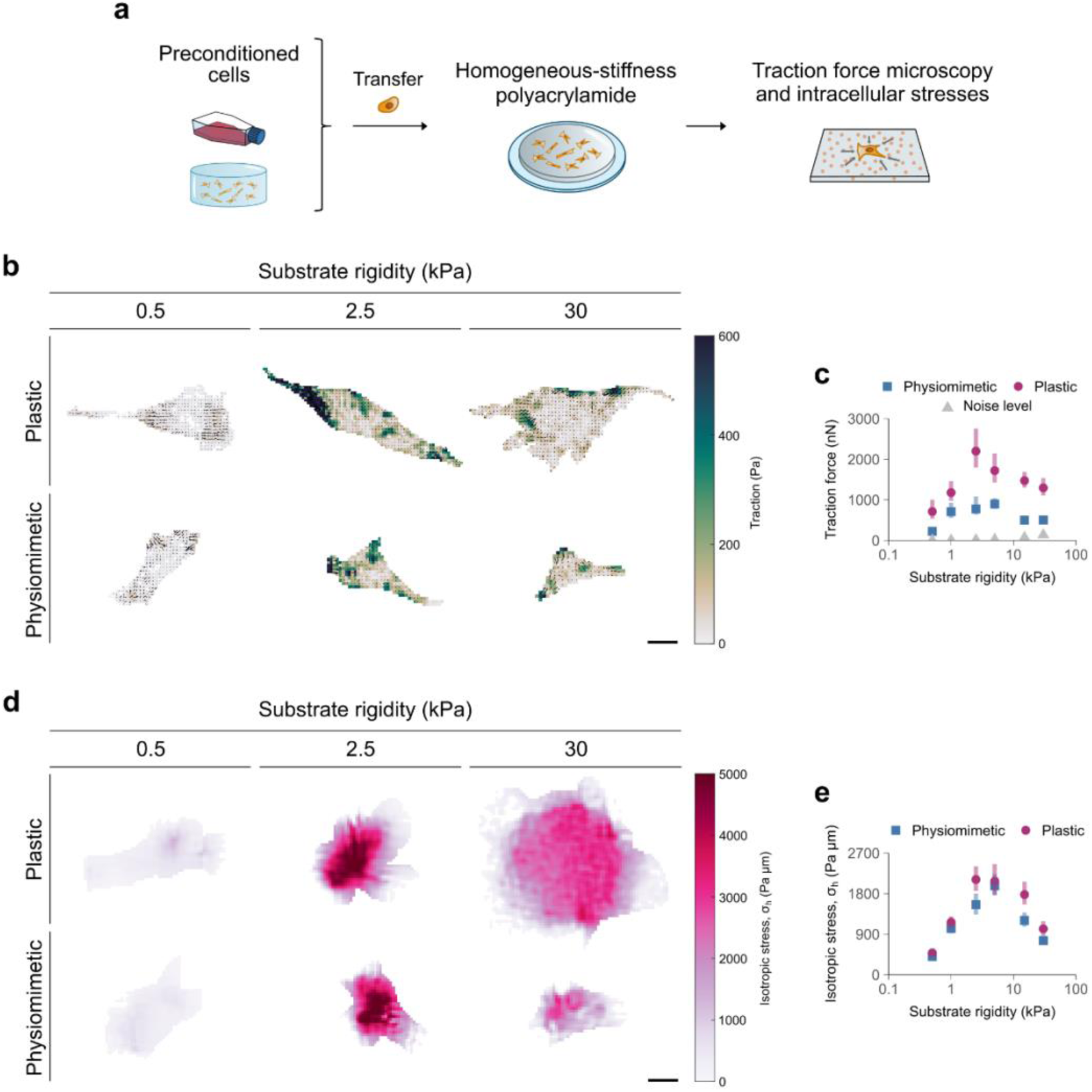
Preconditioning modulates traction force magnitude but preserves biphasic force-stiffness behavior. **a,** Experimental design. RFL-6 fibroblasts were cultured for 4 days on plastic or in physiomimetic hydrogels. After preconditioning, cells were replated on PAA gels (0.5–30 kPa) to quantify traction forces and intracellular stresses. **b,** Representative traction maps of plastic- and physiomimetic-preconditioned fibroblasts cultured on substrates of increasing stiffness (0.5–30 kPa). Arrows indicate traction vectors and the color scale denotes magnitude. Scale bar, 25 μm. **c,** Quantification of cell traction force as a function of substrate stiffness. Both populations displayed biphasic force-stiffness relationships with distinct optimal stiffness (∼2.5 kPa for plastic-preconditioned cells and ∼5 kPa for physiomimetic-preconditioned cells). Across all rigidities, traction magnitudes were 40–70% lower in physiomimetic-preconditioned cells. A permutation-based quadratic ANOVA confirmed a strong overall effect of preconditioning on force output (condition: *p* = 0.0001) and revealed significant contributions of both the linear (*p* = 0.0063) and quadratic (*p* = 0.0027) stiffness terms, consistent with a peak-shaped, biphasic dependence. Noise levels are shown in grey, indicating the minimum detectable force for each substrate stiffness (see Methods). **d,** Representative stress fields reconstructed from traction maps using 2D force-balance equations (see Methods). Cytoskeletal tension corresponds to the isotropic component of the stress tensor. **e,** Mean isotropic stress averaged over the cell spreading area for plastic- and physiomimetic-preconditioned cells across substrate rigidities. A permutation-based ANOVA confirmed that both preconditioning and stiffness showed significant main effects (*p* < 0.0002 and *p* = 0.0222, respectively). Post-hoc permutation tests showed that below 5 kPa, plastic-preconditioned cells tended to generate higher intracellular stresses (∼20% on average), with significant differences at specific stiffness levels. The two conditions displayed similar stress at 5 kPa (*p* = 0.64). At higher rigidities, plastic-preconditioned cells exhibited markedly greater intracellular tension (∼40% on average). Data in panels **(c)** and **(e)** represent mean ± 95% confidence intervals, estimated by bootstrap resampling of 45–51 cells per stiffness and condition from three independent experiments.

To reveal how forces are internally distributed, we reconstructed the in-plane stress field from traction force maps by applying Newton’s laws of force balance^27^. Cytoskeletal tension was quantified by the isotropic component of the stress tensor, averaged over the cell spreading area (σ_h_, Fig. 2d), reflecting isotropic compression or tension within the cytoskeleton transmitted through adhesions. A permutation-based two-way ANOVA revealed a significant effect of preconditioning (*p* < 0.0002) and stiffness (*p* = 0.0222) on mean intracellular stress (Fig. 2e). As for traction forces, isotropic stress showed a biphasic dependence on rigidity, peaking at ∼2.5 kPa for plastic-preconditioned cells and ∼5 kPa for physiomimetic-preconditioned cells. In the low-stiffness regime (<5 kPa), plastic-preconditioned cells generated moderately higher stresses (∼20%), reaching significance at selected rigidities, whereas at 5 kPa levels converged (*p* = 0.64). Above this rigidity, plastic-preconditioned cells maintained significantly higher tension (∼40%). This indicates that plastic preconditioning stabilizes a high-tension state at elevated stiffness. In contrast, physiomimetic-preconditioned cells reduced intracellular tension beyond 5 kPa. Overall, plastic preconditioning promotes intracellular tension at high stiffness, biasing cells toward a high-tension mechanical regime.

### Preconditioning sets durotactic direction

Previous studies have shown that cells displaying a peak in traction forces tend to accumulate at their optimal stiffness^11^. To test this prediction, we used collagen I-coated PAA gels featuring a continuous stiffness gradient spanning ∼0.2 to 20 kPa, a range that extends across both healthy and fibrotic lung tissue^28–30^ (see Methods, Fig. 3a and Extended Data Fig. 3). Cells were seeded uniformly across the gradient and treated with thymidine to arrest proliferation, thereby eliminating potentially confounding effects arising from differential stiffness-dependent cell growth. Phase-contrast images were acquired 4 h after seeding and again after 48 h, and the relative change in local cell density (Δ*ρ*) was quantified in percentage points (pp, see Methods) as a readout of stiffness-dependent cell accumulation (Fig. 3a).

**Figure 3.**
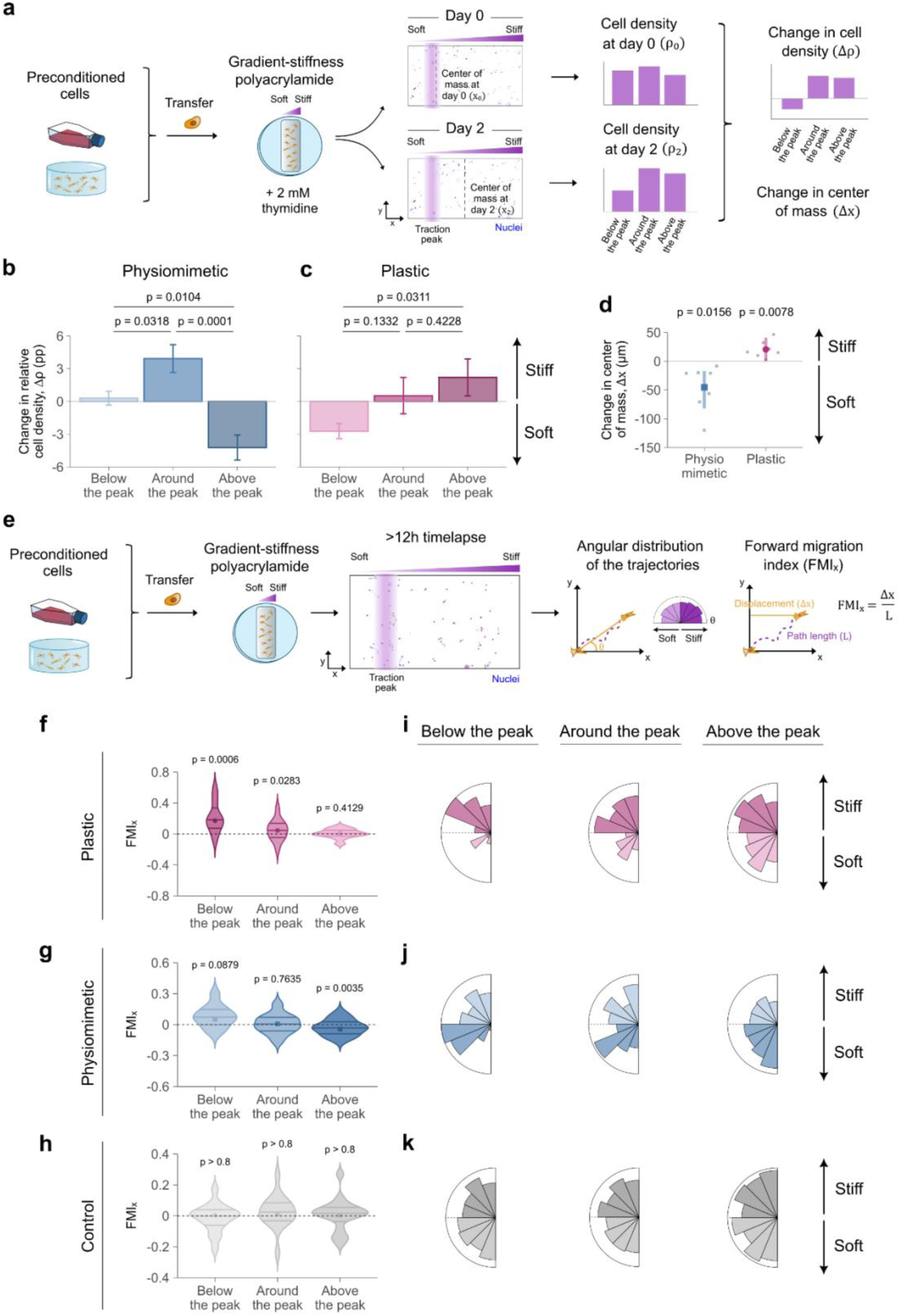
Plastic and physiomimetic preconditioning set opposite durotaxis directions. **a**, Experimental setup for cell accumulation experiments. PAA gels with continuous stiffness gradients (∼0.2**–**20 kPa) were used to assess stiffness-dependent cell accumulation. Uniformly seeded cells were treated with thymidine to arrest proliferation. Relative changes in cell density after 48 h (Δ*ρ*) were quantified in percentage points (pp), together with the change in the center of mass (Δ*x*). **b–c**, Stiffness-dependent cell redistribution. Physiomimetic-preconditioned cells accumulated around their stiffness peak, depleting stiffer regions consistent with negative durotaxis **(b)**. In contrast, plastic-preconditioned cells accumulated at stiffer regions, consistent with positive durotaxis **(c)**. Data are mean ± SEM. *p*-values were obtained by fitting a linear mixed-effects model with gel identity as a random intercept (see Methods). **d,** Center of mass (Δ*x*) net displacement. Negative values indicate migration toward soft (negative durotaxis) and positive values indicate migration toward stiff (positive durotaxis). Data are mean ± 95% confidence intervals estimated by bootstrap resampling of 7 gels per condition from three independent experiments. *p*-values were computed using two-tailed permutation tests comparing the mean Δ*x* of each group and gel against zero. Data on panels **b–d** represent n = 7 gels from three independent experiments with >115 cells per gel. **e,** Experimental setup for time-lapse durotaxis assays. Cells were uniformly seeded on PAA gels with continuous stiffness gradients (∼0.2**–**20 kPa) and tracked over 12 h. Individual cell trajectories were used to compute the forward migration index (*FMI_x_*) and the angular distributions. **f–h,** *FMI_x_* values for plastic, physiomimetic, and control cells measured below, around, and above the optimal stiffness peak. Below the peak, both preconditioned groups exhibited positive *FMI_x_*values consistent with positive durotaxis. Around the peak, plastic-preconditioned cells displayed significant durotaxis, while physiomimetic-preconditioned cells remained unbiased. Above the peak, plastic-preconditioned cells showed a positive but nonsignificant trend, whereas physiomimetic-preconditioned cells exhibited significantly negative *FMI_x_* values, indicative of negative durotaxis. Control cells correspond to plastic-preconditioned cells migrating on uniform stiffness gels matching each stiffness region. n = 31–465 cells per group, from four independent experiments; tests are two-tailed permutation tests comparing mean *FMI_x_* values against zero. **i–k,** Angular distributions of migration trajectories (first minus last position) below, around and above the optimal stiffness peak.

Physiomimetic-preconditioned cells exhibited a pronounced accumulation (Δ*ρ*>0) around their optimal stiffness range (≈600–900 µm; Extended Data Fig. 4a). Below this region (<600 µm), cell density varied only minimally, consistent with a weak positive durotaxis response in softer regions, as previously reported^11^. In contrast, density declined sharply (Δ*ρ*<0) above the peak (>900 µm), indicating that accumulation near the optimum primarily resulted from cells migrating down the stiffness gradient from stiffer regions, consistent with negative durotaxis. Phenomenological simulations reproduced this redistribution pattern for negatively durotactic cells (Supplementary Text II and Extended Data Fig. 5a-c). Strikingly, plastic-preconditioned cells displayed a fundamentally different behavior: rather than depleting stiff regions, they preferentially depleted soft regions (<350 µm) and accumulated beyond their optimal stiffness (>350 µm; Extended Data Fig. 4b), a pattern incompatible with negative durotaxis but consistent with a positive-durotactic regime (Supplementary Text II and Extended Data Fig. 5d).

To accurately assign local stiffness values, each gradient was characterized by atomic force microscopy (AFM) and spatial coordinates were mapped to Young’s modulus, enabling partitioning into regions below, around, and above the peak (see Methods and Fig. 3b-c). Linear mixed-effects analysis confirmed that physiomimetic-preconditioned cells accumulated significantly at the peak relative to both adjacent regions (*p* = 0.0318 and *p* = 0.0001, Fig. 3b), whereas plastic-preconditioned cells showed no significant accumulation at the peak compared with either neighboring region (*p* = 0.1332 and *p* = 0.4228, respectively), but instead exhibited a significant enrichment above the peak relative to stiffnesses below it (*p* = 0.0311, Fig. 3c). Consistent with model predictions (Supplementary Text II and Extended Data Fig. 5i), center-of-mass analysis revealed significant shifts toward softer regions for physiomimetic-preconditioned cells and toward stiffer regions for plastic-preconditioned cells, indicating opposite durotactic biases despite similar traction force optima (Fig. 3d). Taken together, these results suggest that although both populations display a traction force peak, their durotactic responses are opposite in direction: plastic-preconditioned cells undergo positive durotaxis across all stiffnesses tested, whereas physiomimetic-preconditioned cells exhibit a negative-durotactic bias on the stiff side of the traction force peak, leading to accumulation near their optimal stiffness.

To characterize the migration dynamics underlying the observed cell redistribution, we performed time-lapse imaging on substrates of uniform or graded stiffness. Short-term imaging (1 h, 1 s intervals) on uniform 0.5, 5, and 30 kPa gels revealed that plastic-preconditioned cells were more dynamic (especially on 0.5 kPa), with pronounced lamellipodia and larger displacements, whereas physiomimetic-preconditioned cells displayed milder protrusions and limited movement (Extended Data Fig. 6a and Supplementary Video 1). Over 12 h (20 min intervals) on uniform gels ranging 0.5–30 kPa, migration speed decreased modestly with rigidity in both conditions. Permutation-based ANOVA revealed significant effects of stiffness (*p* < 0.0001) and condition (*p* < 0.0001), with plastic-preconditioned cells migrating ∼2.5-fold faster than physiomimetic-preconditioned cells (Extended Data Fig. 6b; Supplementary Video 2). Despite slower speeds, physiomimetic-preconditioned cells displayed greater directional persistence (Extended Data Fig. 6c). Having established the baseline dynamics on uniform substrates, we next examined whether these distinct migratory behaviors gave rise to directional migration biases on stiffness gradients.

We quantified durotaxis from time-lapse imaging on gradients ranging from ∼0.2 to 20 kPa (Fig. 3e) using the forward migration index (*FMI_x_*), which measures net cell displacement parallel to the gradient normalized by the total path length (Fig. 3e, inset). To determine directional bias, we compared the mean *FMI_x_* of each group and stiffness region against zero (null hypothesis of adurotaxis). Below the optimal stiffness peak, both populations exhibited positive durotaxis, reaching strong statistical significance for plastic (*p* = 0.0006; Fig. 3f) and near significance for hydrogels (*p* = 0.0879; Fig. 3g). At stiffness values close to the peak, plastic-preconditioned remained positively durotactic (*p* = 0.0283; Fig. 3f), whereas physiomimetic-preconditioned cells were adurotactic (*p* = 0.7635; Fig. 3g). Above the stiffness peak, plastic-cultured cells showed a positive durotactic trend that did not reach statistical significance (*p* = 0.4129; Fig. 3f), consistent with the attenuation of positive durotaxis at higher substrate rigidities reported previously^8,9,31^. In contrast, physiomimetic-preconditioned cells exhibited a statistically significant negative *FMI_x_* (*p* = 0.0035; Fig. 3g), indicative of negative durotaxis. As expected, control cells within each stiffness region did not display a significant directional trend (*p* > 0.8, all controls; Fig. 3h). Analysis of the angular distribution of migration trajectories further supported these findings, revealing opposite durotactic biases for plastic- and physiomimetic-preconditioned cells above the optimal stiffness, while control cells remained unbiased (Fig. 3i-k). Taken together, these findings indicate that, in parallel with alterations in cytoskeletal organization and traction forces, preconditioning determines durotactic direction: plastic favors migration toward stiffness, whereas physiomimetic favors migration toward softness.

### Molecular clutch identifies regimes of durotaxis reversal

Physiomimetic preconditioning fundamentally altered cytoskeletal architecture, FA organization, and durotactic directionality. To dissect the underlying mechanism, we extended a two-dimensional molecular clutch framework that integrates myosin activity, adhesive clutch bond dynamics, and F-actin assembly^11,19^ (Fig. 4a; Supplementary Text III). In the model, traction forces arise from myosin-driven retrograde actin flow resisted by molecular clutches that couple the cytoskeleton to the substrate. Load-dependent clutch dissociation and motor force generation produced the expected biphasic traction-stiffness relationship^32,33^ (Fig. 4b,c): weak traction at low stiffness due to premature clutch failure, maximal transmission at intermediate stiffness, and reduced coupling at high stiffness due to rapid force buildup and frictional slippage.

**Figure 4.**
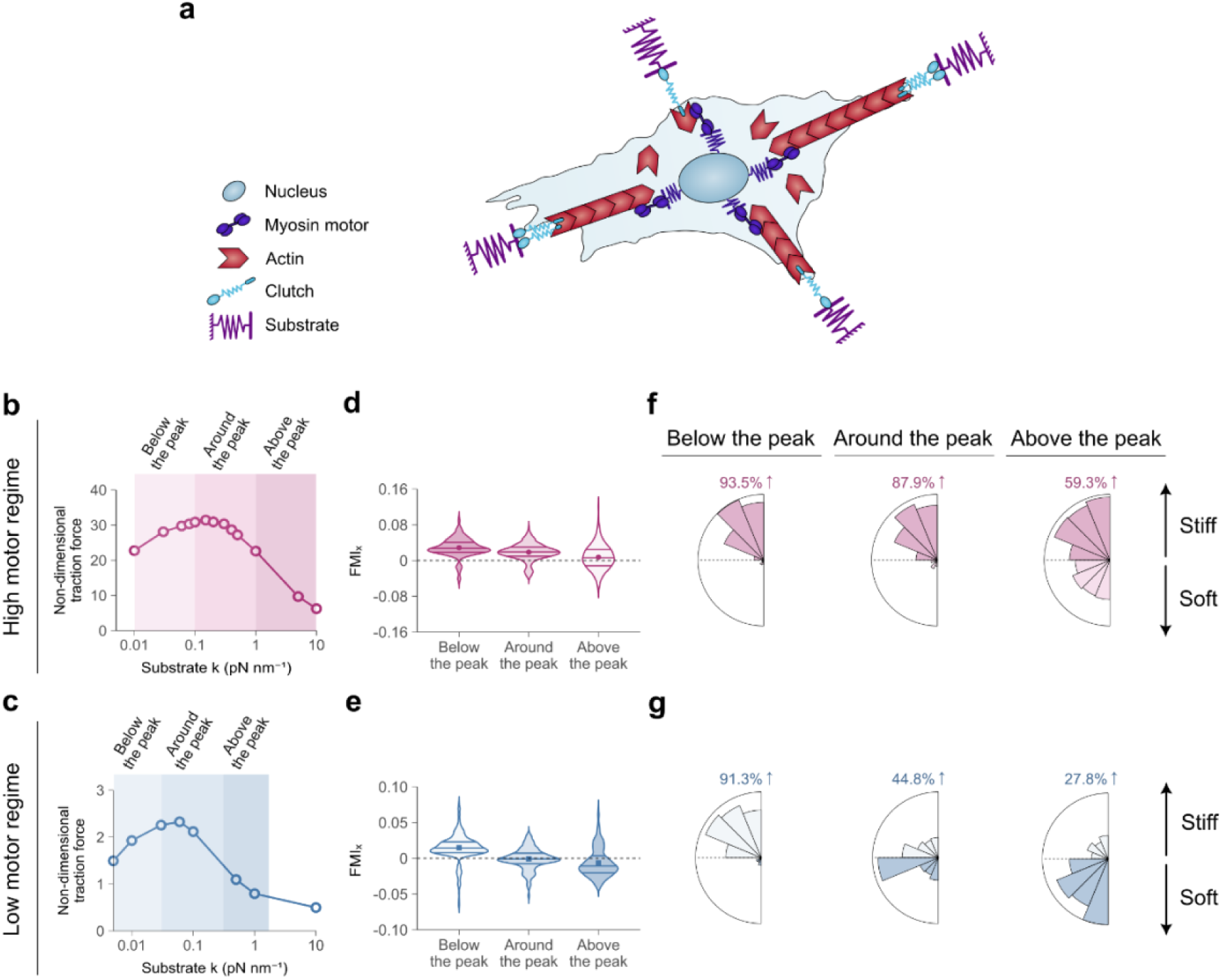
Molecular clutch model predicts distinct durotactic behaviors depending on motor-adhesion regime. **a,** Schematic of the two-dimensional motor-clutch cell migration model. Actin filaments are pulled rearward by myosin motors and transiently coupled to the substrate through molecular clutches. The balance between motor activity, clutch binding dynamics, and substrate stiffness determines traction force generation and cell migration. **b,** Dimensionless total time-averaged traction force as a function of substrate stiffness for the high-motor regime with mechanosensitive motors and adhesion reinforcement. Traction forces exhibit a biphasic dependence on stiffness, defining three stiffness regions: below the peak, around the peak, and above the peak. **c,** Dimensionless traction force as a function of substrate stiffness for the low-motor regime, showing a reduced force magnitude and a shifted biphasic peak compared to the high-motor regime. **d,** Forward migration index (*FMI_x_*) for cells migrating within each region in the high-motor regime. Cells migrate predominantly in the direction of the stiffness gradient across all three regions, consistent with persistent migration toward stiffer substrates, as observed for plastic-preconditioned cells. **e,** *FMI_x_* for the low-motor regime. Migration is weakly biased toward stiffer regions below and around the traction peak, but above the peak cells migrate preferentially toward softer substrates, consistent with experimental observations of physiomimetic-preconditioned cells. **f,** Rose plots showing migration directionality for the high-motor regime below the traction peak, around the peak, and above the peak (n = 400, 380, 400 cells), confirming a bias toward increasing substrate stiffness. **g,** Rose plots of migration direction for the low-motor regime below, around, and above the traction peak (n = 400, 400, 400 cells), revealing a transition from weakly biased migration toward stiffer regions below the peak to preferential migration toward softer regions above the peak. Numbers in pink and blue indicate the percentage of cells whose net migration direction had a positive projection along the stiffness gradient (positive durotaxis). Model parameter values are provided in Extended Data Table 4.

Simulations revealed that the slope of the traction-stiffness curve alone does not determine durotactic directionality. While traction curves reflect time-averaged forces, directional migration emerges from time-dependent competition between protrusions. Plastic-preconditioned cells were modeled in a high-motor, moderately reinforced adhesion regime with mechanosensitive motor recruitment (Supplementary Text III, Eq. 11), consistent with increased phosphorylated myosin II levels on stiff substrates (Fig. 1g-i). This regime is further justified by the phenotypic signature of plastic-preconditioned cells, namely their large traction forces, sustained migration speed at high stiffness, and monotonic increases in spreading and adhesion growth with rigidity (Supplementary Text III, Eq. 12). Because force-loading rates increase with stiffness, protrusions oriented toward stiffer regions accumulate force more rapidly, gaining a temporal advantage during adhesion maturation. Adhesion reinforcement delays clutch failure, allowing stiff-side protrusions to dominate even beyond the tiome averaged traction peak. Simulations predicted robust positive durotaxis below the peak (*FMI_x_*> 0; 93.5% up-gradient migration), moderate positive durotaxis around the peak (*FMI_x_* > 0; 87.9% up-gradient migration), and weak positive durotaxis above the peak (*FMI_x_* > 0; 59.3% up-gradient migration), with loss of directionality only at very high rigidity due to symmetric frictional slippage (Fig. 4d,f), consistent with experiments (Fig. 3f). Notably, a stiffness-insensitive high-motor, high-clutch regime without reinforcement also sustained positive durotaxis (Extended Data Fig. 7), indicating that mechanosensitive feedback is not strictly required when motor and clutch numbers are sufficiently large.

By contrast, physiomimetic preconditioning corresponded to a low-motor, non-mechanosensitive regime, which we modeled using stiffness-insensitive low-motor recruitment in the absence of adhesion reinforcement. In this regime, simulations predicted positive durotaxis below the traction peak (*FMI_x_* > 0; 91.3% up-gradient migration), consistent with preferential stabilization of stiff-oriented protrusions under moderate loading (Fig. 4e,g). Around the peak, directionality weakened substantially (*FMI_x_* ∼ 0; 44.8% up-gradient migration), indicating near-adurotactic behavior (Fig. 4e,g). Above the peak, rapid force accumulation on the stiff side exceeded clutch lifetimes, promoting slippage and destabilizing stiff-side protrusions, resulting in negative durotaxis (*FMI_x_* < 0; 27.8% up-gradient migration) (Fig. 4e,g), consistent with experiments (Fig. 3g). At very high stiffness, symmetric slippage again abolished directional bias.

Together, these simulations demonstrate that negative durotaxis requires not only a traction force optimum, but a mechanical regime in which rapid force accumulation on stiff substrates destabilizes protrusions. Pharmacological reducing myosin II activity is predicted to shift plastic-preconditioned cells into an effectively low-motor, low-clutch regime, lowering loading rates and minimizing adhesion reinforcement. In this regime, stiff-side protrusions are expected to fail before acquiring a temporal advantage, resulting in negative durotaxis. Accordingly, plastic-preconditioned cells are predicted to reverse durotactic directionality upon reduction of myosin II activity, which we tested experimentally.

### Myosin motors inhibition switches durotaxis direction

According to our framework, elevated myosin activity in plastic-preconditioned cells sustains positive durotaxis by stabilizing stiff-side protrusions despite the presence of a traction force peak (Fig. 4b,d). The model predicts that reducing motor contractility shifts cells into an effectively low-motor, low-clutch regime in which stiff-side protrusions fail before outcompeting protrusions on the softer side, thereby preventing stiff-side dominance and promoting negative durotaxis beyond the traction force peak. To test this prediction, we partially inhibited non-muscle myosin II in plastic preconditioned cells using para-nitro-blebbistatin (hereafter blebbistatin). We tested a range of blebbistatin concentrations and selected a dose that induced measurable cytoskeletal alterations without significantly affecting cell migration speed (Extended Data Fig. 8). Among the conditions examined, 10 µM blebbistatin produced pronounced cytoskeletal changes while maintaining migration speeds comparable to untreated cells (Extended Data Fig. 8).

Blebbistatin-treated cells attenuated their sensitivity to substrate stiffness (Fig. 5a). Cell spreading area initially increased with stiffness at a rate comparable to DMSO-treated controls, but sharply decreased on stiff substrates (15 and 30 kPa; Fig. 5b). FAs increased in length with stiffness at a similar pace to controls, yet remained significantly shorter across all rigidities (Fig. 5c). Adhesions were broadly distributed throughout the cell body with only weak peripheral enrichment, contrasting the pronounced peripheral localization observed in DMSO-treated controls (Extended Data Fig. 9). The actin cytoskeleton appeared thin and disorganized, and lacked preferential alignment (Fig. 5a,d). Blebbistatin treatment reduced traction forces by ∼40% and shifted the stiffness optimum from 2.5 to 5 kPa (Fig. 5e), in line with that observed for physiomimetic-preconditioned cells (Fig. 3c). Together, these results indicate that partial inhibition of myosin activity diminishes mechanosensitivity, effectively recapitulating a mechanical phenotype similar to physiomimetic-preconditioned cells.

**Figure 5.**
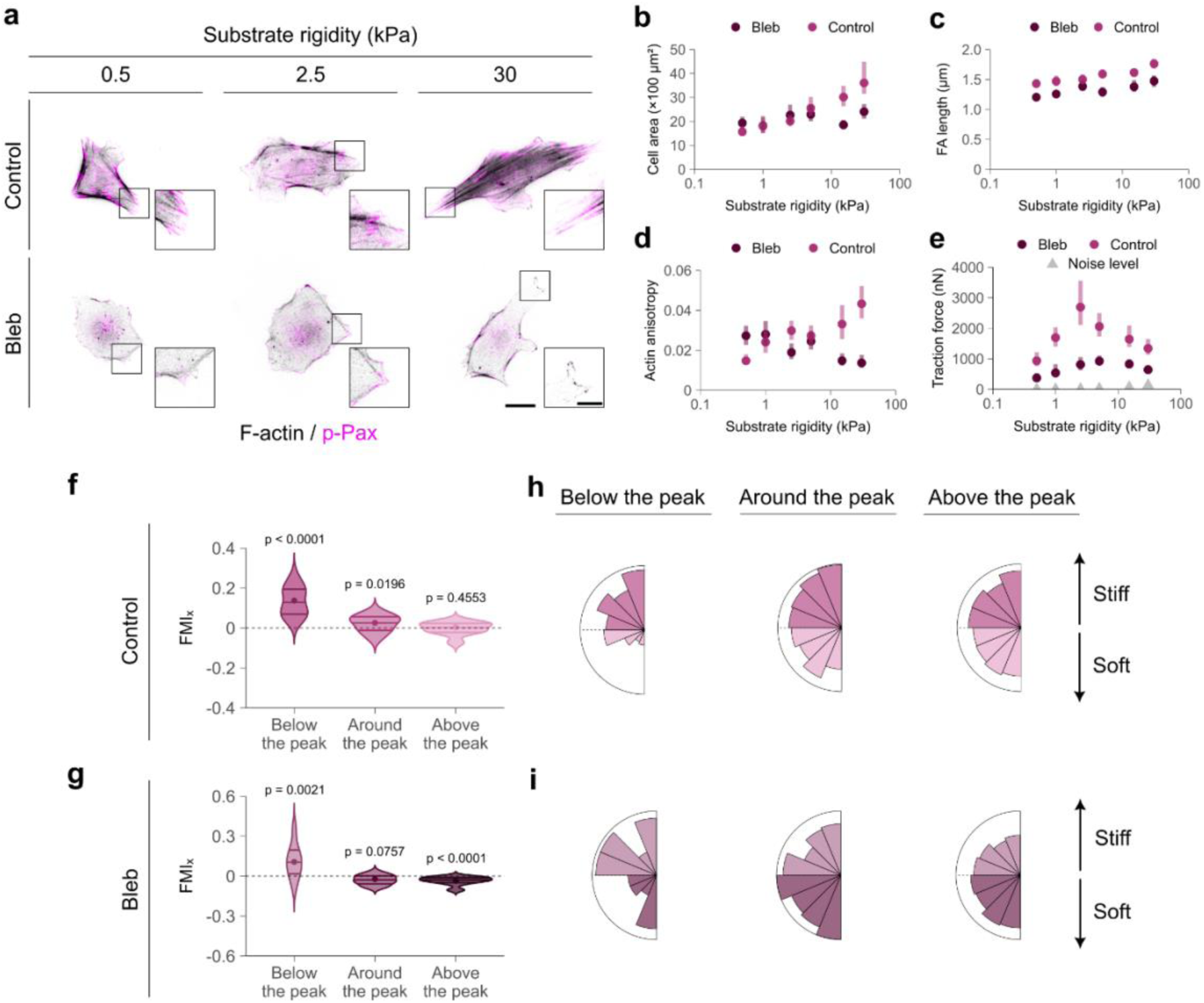
Inhibition of myosin motors reverses positive durotaxis in plastic-preconditioned cells. **a,** Representative images of actin filaments (phalloidin) and focal adhesions (FAs; p-Paxillin) in plastic-preconditioned cells treated with DMSO (Control) or 10 µM blebbistatin (Bleb) on 0.5, 2.5, and 30 kPa substrates. Scale bar, 25 µm. Inset, magnified view highlighting individual FAs (scale bar, 10 µm). **b,** Cell spreading area across various substrate stiffness. Control cells spread more extensively, displaying rigidity-dependent growth, whereas blebbistatin-treated cells showed a similar trend below 5 kPa, but at stiffer regions cell growth diminished. **c,** FA length as a function of substrate stiffness. FA length increased with substrate stiffness in both control and blebbistatin-treated cells. Control cells consistently exhibited larger FAs. **d,** Quantification of actin fiber alignment. Control cells displayed increased anisotropy at stiffer rigidities, while blebbistatin-treated cells lacked preferential alignment. Permutation-based two-way ANOVA revealed significant treatment effects in panels **b-d** and stiffness-dependent responses for spreading area and FA length, with a significant condition×stiffness interaction for anisotropy (all *p*≤ 0.0351). **e,** Cell traction force as a function of substrate stiffness. Both populations displayed biphasic force-stiffness relationships with distinct optimal stiffness (∼2.5 kPa for control cells and ∼5 kPa for blebbistatin-treated ones). Permutation-based quadratic ANOVA showed a strong effect of treatment (*p* = 0.0001) and significant linear (*p* = 0.028) and quadratic (*p* = 0.018) stiffness terms, indicating a biphasic force-stiffness relationship in both populations. Noise levels are shown in grey, indicating the minimum detectable force for each substrate stiffness (see Methods). Error bars in panels **b–e** indicate mean ± 95% confidence intervals computed by bootstrapping resampling of 30–57 cells per stiffness and condition pooled from three independent experiments. **f–g,** Forward migration index (*FMI_x_*) values for control and blebbistatin-treated cells. Below the peak, both groups exhibited significantly positive *FMI_x_*values, consistent with positive durotaxis. Around the peak, control cells displayed significant durotaxis, while blebbistatin-treated cells showed a non-significant bias towards negative durotaxis. Above the peak, control cells exhibited a non-significant positive trend, while blebbistatin-treated cells showed significantly negative *FMI_x_* values, indicating negative durotaxis (n = 77–1570 cells per group from three independent experiments; two-tailed permutation tests). **h–i,** Angular distributions of migration trajectories (first minus last position) below, around and above the optimal stiffness peak.

To test whether reducing force generation in plastic-preconditioned cells induces negative durotaxis, we compared blebbistatin-treated cells with DMSO-treated controls migrating on stiffness gradients and quantified directional bias using the *FMI_x_* along the gradient. As in previous experiments, directional bias was measured using two-sided one-sample permutation tests (10,000 resamples) comparing mean *FMI_x_* values against zero. Cells were grouped according to whether they migrated in regions below, near or above their optimal stiffness. Control cells exhibited pronounced positive durotaxis below the peak (*p* < 0.0001). This bias weakened near the peak but remained significantly positive (*p* = 0.0196), and became positive but non-significant above the peak (*p* = 0.4553; Fig. 5f), consistent with the behavior of plastic-preconditioned cells (Fig. 3f). By contrast, blebbistatin-treated cells exhibited positive durotaxis below the peak (*p* = 0.0021), became negative but non-significant near the peak (*p* = 0.0757) and shifted to a pronounced, and statistically significant negative durotaxis above the peak (*p* < 0.0001; Fig. 5g), closely resembling the response of physiomimetic-preconditioned cells (Fig. 3g). The angular distribution of migration trajectories independently corroborated this reversal, revealing a clear bias toward migration down stiffness gradients in the presence of blebbistatin (Fig. 5h,i). Together, these data demonstrate that the presence of a traction force peak alone is insufficient to drive negative durotaxis. Rather in the cells examined here, negative durotaxis emerges only when cells operate within a low motor activity mechanical regime.

## Discussion

Durotaxis is increasingly recognized as a key mechanism driving directed cell migration in development^10^ and disease^35^. While most cells migrate toward stiffer environments through positive durotaxis^4,36^, recent studies have shown that this response is not universal, and that certain cells instead migrate toward softer regions through negative durotaxis^37,11^. Here we show that culture conditions alone, in the absence of genetic or biochemical manipulation, are sufficient to tune durotaxis direction. We cultured fetal rat lung fibroblasts in either plastic or a 3D physiomimetic hydrogel derived from decellularized porcine lungs. Although both plastic- and physiomimetic-conditioned cells exhibited a biphasic traction force-stiffness relationship with a maximum at intermediate rigidity, their migratory responses were strikingly different. Plastic-cultured cells exhibited elevated traction forces and migrated toward stiffer regions, exhibiting positive durotaxis. In contrast, brief preconditioning of fibroblasts within physiomimetic hydrogels for only a few days was sufficient to reduce overall force magnitude and reverse durotactic behavior, driving migration toward softer regions and accumulation near physiological lung stiffness (∼5 kPa). We developed a molecular clutch model that captures the phenomenology and shows that a biphasic force-stiffness relationship is necessary but not sufficient to generate negative durotaxis.

Positive durotaxis has long been regarded as the canonical form of stiffness-guided migration in fibroblasts, as first identified by Lo and colleagues^3^ more than 20 years ago. Subsequent studies across diverse mesenchymal^6,7,9^, epithelial^8^ and even amoeboid^38^ cell types have consistently reproduced this behavior, establishing positive durotaxis as a characteristic signature of cellular mechanosensitivity. More recently, this canonical response has been challenged, but almost exclusively in specialized cell types. For example, a subpopulation of weakly adherent human breast cancer MDA-MB-231 cells shows little or no sensitivity to stiffness gradients^39^, whereas downregulation of talin in the same cell line induces migration toward softer regions^11^. Negative durotaxis has also been observed in developmental contexts, such as during formation of the *Xenopus* optic pathway, where retinal ganglion cell axons extend toward softer, more compliant tissue^37^. Likewise, human glioblastoma U251-MG cells migrate preferentially toward softer regions, accumulating at an optimal stiffness of ∼8 kPa and exhibiting pronounced negative durotaxis. Our results extend these observations by identifying fetal rat lung fibroblasts (RFL-6) as a new system capable of exhibiting negative durotaxis under mechanical regimes established by physiomimetic preconditioning or myosin inhibition.

The combined experimental and modeling results show that the transition between positive and negative durotaxis is set by the mechanical regime of the coupled cytoskeleton-adhesion system, rather than by the slope of the traction-stiffness relationship alone, as previously proposed^11^. Although traction forces generally increase monotonically with substrate rigidity^21–26^, fetal fibroblasts exhibit a biphasic traction-stiffness response, characterized by a pronounced maximum at intermediate stiffness. Importantly, the presence of a traction force optimum is not sufficient to predict durotactic directionality. Cells preconditioned on plastic migrate persistently up stiffness gradients even beyond the force maximum, consistent with a high-motor, moderately reinforced adhesion regime in which rapid force loading on stiff substrates, combined with delayed clutch failure, confers a temporal advantage to stiff-side protrusions. By contrast, physiomimetic preconditioning places cells in a low-motor, weakly reinforced adhesion regime, where rapid force buildup on stiff substrates outpaces clutch engagement, destabilizes stiff-side protrusions, and allows softer-side protrusions to dominate, resulting in negative durotaxis. Durotaxis has often been explained by assuming that cells migrate toward regions where they generate larger traction forces, implicitly linking directionality to time-averaged force-stiffness relationships. This work instead indicates that durotactic direction emerges from the dynamic competition between protrusions, offering a more mechanistic understanding of the phenomenon.

In line with this framework, the model predicts that reducing myosin II activity in plastic-preconditioned cells destabilizes adhesions at sufficiently high stiffness and induces negative durotaxis. Consistent with this prediction, partial inhibition of myosin II was sufficient to induce negative durotaxis in plastic-cultured fibroblasts, supporting the notion that durotactic direction is set by the mechanical regime in which cells operate. Notably, this behavior contrasts with observations in glioblastoma U-251-MG cells, where reduced myosin II motor activity converted negative durotaxis into positive durotaxis^11^. In that system, reduced contractility shifted the optimal stiffness three-fold, from ∼8 to ∼24 kPa, causing cells to accumulate at the stiff end of the gradient. In contrast, the peak shift observed here was modest (from ∼2.5 to ∼5 kPa), and negative durotaxis emerged beyond the traction force peak, indicating that reduced contractility can produce negative durotaxis through distinct mechanical landscapes. Together, these findings indicate that the influence of contractility on durotaxis is intrinsically context-dependent, producing different outcomes depending on the regime at which cells operate.

An important open question is to what extent these findings apply to other cell lines. Here, we focused on fetal rat lung fibroblasts, that although they are not strictly primary, they are non-transformed and may retain features that allow them to show negative durotaxis. By contrast, more transformed fibroblast lines, such as MEFs or NIH3T3 cells, typically exhibit elevated contractility and strong adhesion reinforcement^40,41^, properties commonly associated with robust positive durotaxis, and may consequently be less responsive to physiomimetic preconditioning or unable to show negative durotaxis. Importantly, the mechanisms by which 3D preconditioning efficiently induces a negative durotaxis phenotype remain unknown. This effect may result from a combination of factors, including the 3D architecture, matrix stiffness, collagen presentation, and tissue-specific biochemical signals, including matrikine fragments. Supporting this idea, 3D preconditioning in collagen matrices has been shown to induce a rejuvenated state in fibroblasts that differs from that obtained by 2D stiffness preconditioning alone^42^. This suggests that dimensionality may engage mechanobiological programs that cannot be recapitulated by substrate stiffness alone. Future studies will be needed to identify the minimal physical and biochemical cues required for this switch and to understand how they place cells in a low-sensitivity state that allows durotaxis reversal.

Our findings suggest that physiological-like conditions bias durotaxis towards soft environments. Given that cells *in vivo* reside within compliant tissues, we speculate that our preconditioning likely restores a fundamental physiological migratory program. In contrast, prolonged culture on rigid substrates promotes a hypercontractile phenotype reminiscent of fibrotic or tumor-associated stroma. As most cell lines used in durotaxis studies are routinely maintained on plastic, stiff culture conditions may explain why most cell lines display robust positive durotaxis^3,6–9,31^. This framework has broad implications for understanding stiffness-guided migration in health and disease. Physiomimetic-preconditioned cells exhibited negative durotaxis and accumulated near ∼5 kPa, corresponding to the physiological stiffness of lung tissue^28,29,43^. This suggests that stromal cells may preferentially migrate toward the stiffness of their native environment, contributing to mechanical homeostasis. In pathological contexts such as fibrosis or cancer, where tissue stiffness increases, this preference may shift, favoring positive durotaxis and accumulation in abnormally stiff regions^35,44^. Indeed, positive durotaxis has been implicated in myofibroblast enrichment within fibrotic tissue, thereby amplifying disease progression. Pharmacological inhibition of the FAK-paxillin interaction selectively blocks positive durotaxis and reduces fibrosis *in vivo*^35^. Together, our results support a framework in which negative durotaxis characterizes healthy tissue, whereas positive durotaxis emerges as a mechanical hallmark of pathological stiffening.

### Conclusions

Our findings reveal that the direction of stiffness-guided migration is not an immutable cellular property, but a switchable and adaptive behavior encoded by culture conditions and cytoskeletal state. Brief preconditioning of fetal lung fibroblasts within physiomimetic 3D scaffolds was sufficient to shift cells from a high-tension to a low-tension motor-clutch regime, reversing the canonical direction of durotaxis. Molecular clutch simulations showed that while a traction force peak is required for negative durotaxis, it is not sufficient: directionality also depends on the mechanical regime in which cells operate, where reduced contractility lowers force-loading rates on both sides, yet force still accumulates faster on the stiff side, causing stiff-side protrusions to fail before they can win the competition and thereby biasing migration toward softer regions. Physiomimetic preconditioning thus emerges as a powerful strategy to restore cellular mechanical phenotype and recover physiological responses to stiffness cues. More broadly, the ability to reprogram durotactic directionality offers potential routes to therapeutically steer cell migration, for example by directing fibroblasts away from fibrotic lesions or reshaping stromal-tumor interactions. Ultimately, these findings suggest a framework in which durotaxis emerges as a key mechanism for mechanical homeostasis, whereby cells navigate toward environments that align with their internal contractile setpoint.

## Methods

### Cell culture

Rat fetal lung fibroblasts (RFL-6, ATCC; non-transformed line derived from Sprague-Dawley rats) were maintained on standard tissue-culture polystyrene dishes (Grynia) in Kaighn’s Modification of Ham’s F-12 Medium (F-12K, ATCC) supplemented with 20% fetal bovine serum (FBS, Labclinics) and 2% penicillin/streptomycin (Thermo Fisher). Cultures were incubated at 37 °C in a humidified atmosphere containing 5% CO₂. Culture medium was replaced every 2 days following a rinse with 1× phosphate-buffered saline (PBS, Thermo Fisher). Cells were routinely maintained in 75 cm² flasks (Grynia) and passaged every 3-4 days upon reaching approximately 70% confluence. For passaging, cultures were washed with 1× PBS and incubated with 0.25% trypsin-EDTA (Thermo Fisher) for 5 min at 37 °C. Trypsinization was halted by addition of complete medium, and cells were centrifuged at 125 × g for 7 min. The resulting pellet was resuspended in fresh medium and reseeded at appropriate dilutions to allow passaging every 3-4 days.

### Fabrication of ECM-derived physiomimetic hydrogels

Lung ECM hydrogels were prepared as previously described by Pouliot and Falcones^20,45^. Porcine (*Sus scrofa domesticus*) lungs with intact tracheae were obtained from four animals at a local slaughterhouse. Lungs were inspected for structural integrity, and any damaged areas or residual connective tissue were removed. The left lung was converted into a closed perfusion system by clamping the right main bronchus and associated vasculature. Decellularization was performed by sequential perfusion through both the trachea and vasculature with 0.1% Triton X-100 (Sigma-Aldrich) and 2% sodium deoxycholate (Sigma-Aldrich) in ultrapure water for 24 h each at 4 °C, 1 M sodium chloride (Sigma-Aldrich) in ultrapure water for 1 h at 4 °C, and 0.03 mg mL⁻¹ deoxyribonuclease I (Sigma-Aldrich) in distilled water for 1 h at 4 °C. Three rinses with ultrapure water were performed between detergents, and three final rinses with 1× PBS were carried out to remove residual reagents and cellular debris. To assess the decellularization efficiency, genomic DNA was extracted using the PureLink Genomic DNA Mini Kit (Thermo Fisher) according to the manufacturer’s protocol and quantified by spectrophotometry. DNA concentrations were below the accepted threshold of 50 ng mg⁻¹. The decellularized tissue was cut into 1–3 cm² sections excluding cartilaginous regions, frozen at −80 °C for 24 h, freeze-dried (Telsar Lyoquest-55 Plus), and cryomilled in liquid nitrogen (SPEX SamplePrep) to obtain a fine powder. This ECM powder was resuspended at 20 mg mL⁻¹ in 0.01 M hydrochloric acid (VWR Chemicals) and solubilized by digestion with 2 mg mL⁻¹ pepsin from porcine gastric mucosa (Sigma-Aldrich) under continuous stirring at 400 rpm for 16 h at room temperature (RT). The solubilized ECM was diluted 1:9 in 10× PBS and neutralized to pH 7.4 with 0.1 M and 0.2 M sodium hydroxide (Sigma-Aldrich) to obtain a solution ready for cell incorporation.

### Preconditioning in physiomimetic hydrogels or on plastic

Cells were washed once with 1× PBS and incubated with 0.25% trypsin-EDTA for 5 min at 37 °C. Trypsin was neutralized with complete culture medium, and the suspension was centrifuged at 125 × g for 7 min to obtain a pellet. The pellet was resuspended in fresh medium and either seeded on 25 cm² tissue culture flasks (plastic preconditioning; 1.2 × 10³ cells cm⁻²; Grynia) or embedded in ECM hydrogels (physiomimetic preconditioning; 3 × 10⁵ cells mL⁻¹). Hydrogels were then polymerized by incubating at 37 °C for 30 min in 24-well plates (Sigma-Aldrich). Both conditions were maintained under standard culture conditions for 4 days with medium replacement every 2 days.

After preconditioning, cells were recovered by sequential enzymatic digestion. Hydrogels were incubated with 0.25% trypsin-EDTA for 5 min at 37 °C, followed by treatment with 350 U mL⁻¹ collagenase (Gibco) in serum-free medium containing 10% HEPES (Gibco) for 7 min at 37 °C. The digests were gently pipetted to ensure complete dissociation, neutralized with complete medium, and filtered through a 100 µm cell strainer (Sigma-Aldrich) to remove residual matrix fragments. The filtrate was centrifuged at 800 × g for 10 min, and the obtained cell pellet was resuspended in complete culture medium. To maintain consistency across conditions, cells cultured on plastic were subjected to the same recovery protocol, including collagenase treatment, neutralization, and filtration. For experiments, preconditioned cells were seeded on PAA gels at a density of 1.3 × 10³ cells cm⁻² and allowed to adhere for 4 h.

### Preparation of polyacrylamide gels

PAA gels with elastic moduli of 0.5 to 30 kPa were prepared by polymerizing defined mixtures of acrylamide (40%) and bis-acrylamide (2%) (Bio-Rad) in ultrapure water (see Supplementary text I, Extended Data Fig. 1 and Extended Data Table 1). Glass-bottom dishes (35 mm, MatTek) and 12-well glass-bottom plates (Cellvis) were silanized with a Bind-Silane solution (80:2:1 ethanol/acetic acid/Bind-Silane, v/v/v; Sigma-Aldrich) for 5 min to promote gel adhesion. The dishes and plates were then dried with Kimtech wipes (Kimberly-Clark). To prevent adhesion to the top coverslips, 18 mm glass coverslips were immersed for 5 min in Repel-Silane (Cytiva), then cleaned in the same manner. Filtered (0.22 µm) PAA stock solutions were stored at 4 °C in the dark until use. For polymerization, 1 mL of the desired stock was mixed with 0.5% ammonium persulfate (10% w/v in distilled water; Bio-Rad) and 0.05% N,N,N′,N′-tetramethylethylenediamine (Sigma-Aldrich). When required for traction force microscopy, orange fluorescent FluoSpheres polystyrene microspheres (0.2 µm, Thermo Fisher) were added (0.2%). A 20 µL drop of the mixture was placed on a Bind-Silanized substrate and covered with a Repel-Silane-treated coverslip to ensure a flat surface and minimize oxygen inhibition. Polymerization proceeded for 1 h at RT. Gels were then immersed in 10× PBS to facilitate removal of the top coverslip and stored in 1× PBS at 4 °C for up to 7 days.

### Preparation of stiffness gradient polyacrylamide gels

PAA gels with stiffness gradients were fabricated with the sliding mask technique^8,31,46^ (Extended Data Fig. 3). In short, acrylamide (40%) and bis-acrylamide (2%) solutions were mixed with 10× PBS and ultrapure water in defined proportions to generate a stock solution (see Extended Data Table 2). The filtered (0.22 µm) solution was stored at 4 °C in the dark for later. A 5% (w/v) solution of Irgacure 2959 (BASF) in 70% ethanol was used as photoinitiator. For gel polymerization, 985 µL of the acrylamide/bis-acrylamide stock was mixed with 15 µL of Irgacure in an opaque microcentrifuge tube. A 20 µL drop of the mixture was then dispensed onto a Bind-Silanized 35 mm MatTek glass-bottom dish and covered with a Repel-Silane-treated 18 mm glass coverslip to create a flat surface and prevent oxygen inhibition. The assembled sandwich was positioned 10 cm below a 365 nm UV lamp (UVP) and exposed for 245 s using a custom gradient generator that moved an opaque mask across the sample at a constant speed of 10 µm s⁻¹. The progressive exposure created stiffness gradients along the gel through spatial variations in polymerization. After polymerization, gels were immersed in 10× PBS to facilitate removal of the top coverslip, then stored in 1× PBS at 4 °C for up to 7 days.

### Functionalization of polyacrylamide gels

PAA gels were functionalized as previously described^47^. Gels were sterilized in 70% ethanol for 30 min, rinsed three times with 1× PBS, and coated with L-DOPA to enable protein attachment. L-DOPA (2 mg mL⁻¹, Sigma-Aldrich) was prepared in 10 mM Tris buffer (pH 10) at 4 °C for 30 min in the dark, sterilized through a 0.22 µm filter, and applied to the gels for 30 min at RT. After three 1× PBS washes, gels were incubated with rat tail collagen type I (80 µg mL⁻¹, Sigma-Aldrich) in 1× PBS for at least 2 h at 37 °C. Following incubation, gels were washed three times with 1× PBS and stored in 1× PBS at 4°C until use.

### Atomic force microscopy

Uniform and stiffness gradient PAA gels were characterized using a custom-built AFM attached to an inverted optical microscope (Ti-Eclipse, Nikon) as described previously^48^. A pre-calibrated V-shaped cantilever equipped with a 5 µm diameter borosilicate spherical bead and a nominal spring constant of 0.03 N m⁻¹ (Novascan) was used. Deflection sensitivity was obtained from the slope of the force-displacement (*F–z*) curve acquired on a bare coverslip region. For each sample, five *F*–*z* curves (*F* = *k·d*, with *d* the cantilever deflection and *z* the piezotranslator position) were recorded by ramping forward and backward at constant speed (5 µm amplitude, 1 Hz; ∼1 µm indentation). The Young’s modulus (*E*) was obtained by least-squares fitting to the spherical Hertz model:

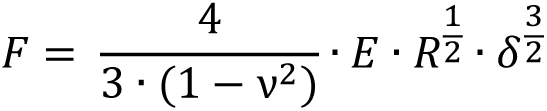

where *ν* is the Poisson’s ratio (assumed 0.5), *R* is the bead radius, and *δ* = *z − z₀ − d* is the indentation depth, with *z₀* representing the contact point. Model fitting was performed using custom MATLAB R2023a (MathWorks, MA) routines.

For uniform gels, three random regions >1 mm apart were selected, with three measurement points spaced ≥20 µm within each region (nine points per gel). For gradient gels, the first measurement point was positioned 100 µm from the softest edge, and subsequent points were acquired at intervals of 100 µm extending toward the stiffest edge.

### Immunofluorescence

Cells were fixed with 4% paraformaldehyde (Sigma-Aldrich) in 1× PBS for 10 min at RT, then washed three times with 1× PBS. Permeabilization was performed with 0.2% Triton X-100 in 1× PBS for 10 min at RT, followed by three 1× PBS washes. Nonspecific binding was blocked with 10% FBS in 1× PBS (blocking buffer) for 30 min at RT. Samples were incubated with primary antibodies diluted in blocking buffer for 1 h at 37 °C: rabbit anti-phospho-paxillin (1:500; Cell Signaling) and mouse anti-phospho-myosin light chain (1:500; Cell Signaling). After three washes with blocking buffer, samples were incubated with corresponding secondary antibodies for 1 h at 37 °C: goat anti-rabbit Alexa Fluor 488 (1:200; Abcam) and goat anti-mouse Alexa Fluor 488 (1:200; Abcam). Following three washes with 1× PBS, conjugated phalloidin-iFluor 555 (1:500; Abcam) was applied when required and incubated for 45 min at RT. Nuclei were stained with Hoechst 33342 (1 drop mL⁻¹; NucBlue, Invitrogen) in 1× PBS for 5 min at RT, followed by three final 1× PBS washes. To minimize photobleaching, samples were mounted face-down on rectangular glass coverslips with Fluoromount-G mounting medium (Invitrogen). Images were acquired on an inverted Nikon Ti microscope equipped with an ORCA-Fusion digital CMOS camera (Hamamatsu) and controlled with MicroManager software^49^ with a CFI Plan Apochromat Lambda D 40×/0.95 NA objective (Nikon).

### Traction force microscopy

Traction force microscopy was performed on an inverted Nikon Ti microscope equipped with an ORCA-Fusion digital CMOS camera (Hamamatsu) and temperature (37 °C), CO₂ (5%), and humidity control using MicroManager software. Images were acquired with a CFI Plan Apochromat Lambda 20×/0.75 NA objective (Nikon), using the phase-contrast channel to visualize cells and the fluorescence channel to image the embedded beads within the PAA gels. For each experiment, two sets of images were recorded: before and after cell detachment, corresponding to the deformed and relaxed states of the substrate, respectively. Cell detachment was achieved by incubating the gels with 10% sodium dodecyl sulfate (Sigma-Aldrich) in distilled water until complete cell removal. Image drift was corrected using the StackReg plugin in Fiji^50^. Bead displacement fields were computed with custom MATLAB software and traction stress maps were obtained by solving the corresponding inverse problem as detailed elsewhere^26^. To estimate the minimum detectable force (noise level) for each substrate stiffness, the same procedure was followed using cell-free gel areas.

### Stress reconstruction from traction maps

In-plane stress fields were reconstructed from traction force microscopy data by using Monolayer Stress Microscopy (MSM)^27^, where 2D force-balance is derived from Newton’s laws. The intracellular stress tensor was computed by applying the inverse stress-recovery formulation, generating spatial maps of *σ_xx_*, *σ_yy_*, and *σ_xy_*. Principal stresses (*σ*_1_, *σ*_2_) were obtained by diagonalizing the stress tensor at each pixel, and the isotropic stress was defined as the isotropic component of the tensor, *σ_h_* = ½(*σ*_1_ + *σ*_2_). For each cell, *σ_h_* was averaged over the segmented cell area to obtain a single scalar measure of cytoskeletal tension. This quantity reflects the local isotropic compression or tension generated within the cytoskeleton and transmitted to the substrate. MSM was implemented in Python 3 using standard libraries.

### Cell accumulation experiments

To evaluate stiffness-dependent cell accumulation, preconditioned cells were seeded at low density on PAA gels containing stiffness gradients (⁓0.2 to 20 kPa). To minimize proliferation-related bias, cell cycle was arrested with thymidine (2 mM; Sigma-Aldrich) 24 h prior to seeding. Nuclei were stained with NucBlue Hoechst 33342 (1 µg mL⁻¹), and identical fields of view were acquired 4 h after seeding (Day 0) and after 48 h (Day 2). Images were acquired on an inverted Nikon Ti microscope equipped with an ORCA-Fusion digital CMOS camera (Hamamatsu) and controlled with MicroManager software with a CFI Plan Fluor 10×/0.30 NA objective (Nikon). For each stiffness gradient, three overlapping images were acquired to span the full gradient range. After imaging, cells were detached with 0.25% trypsin-EDTA (5 min, 37 °C) for AFM mapping of local stiffness.

The overlapping images were stitched using the Grid/Collection Stitching plugin in Fiji. To achieve precise subpixel alignment between time points, Day 0 and Day 2 images were co-registered using the Landmark Correspondences plugin in Fiji. Nuclear centroids were detected using the TrackMate plugin^51^ (Fiji). Cell densities at Day 0 and Day 2 and their difference normalized to the AFM-derived stiffness profile were calculated with custom MATLAB scripts. The center of mass was computed as the mean *x*-coordinate of all nuclei within each gradient. The shift in the center of mass was defined as the difference between Day 2 and Day 0 values.

### Time-lapse imaging

Time-lapse imaging on uniform and gradient PAA gels was performed using an inverted microscope (Eclipse Ti, Nikon) equipped with an ORCA-Fusion digital CMOS camera (Hamamatsu) and controlled with MicroManager software. The microscope was maintained at 37 °C in a humidified atmosphere containing 5% CO₂. Nuclei were stained with NucBlue Hoechst 33342 (1 µg mL⁻¹). Images were acquired with a CFI Plan Fluor 10×/0.30 NA objective (Nikon) in phase-contrast and DAPI channels to visualize cells and nuclei, respectively. The acquisition interval was set to 20 min, and experiments typically lasted >12 h. For migration assays, three random positions >1 mm apart were imaged on gels of uniform stiffness. For durotaxis experiments, three partially overlapping fields spanning the entire stiffness range were imaged.

### Drug treatment

Prior to drug treatment, cells were washed once with 1× PBS and incubated in F-12K medium supplemented with 20% FBS and 2% penicillin-streptomycin. Para-nitro-blebbistatin (10 µM; Motorpharma) was added 1 h before the beginning of the experiment to inhibit myosin II ATPase activity. Dimethyl sulfoxide (DMSO; Sigma-Aldrich) was used as a vehicle control.

### Processing of immunofluorescence images

Fluorescence images were analyzed using Fiji and MATLAB. Nuclei and whole-cell masks were first generated by manual segmentation in Fiji and saved as binary images. A custom MATLAB pipeline was then used to compute cell area, mean p-Myosin intensity and total p-Myosin intensity. Actin anisotropy was quantified using the FibrilTool plugin (Fiji) as described by Boudaoud et al^52^. FAs were segmented using the Focal Adhesion Analysis Server (FAAS)^53^, from which the FA length and density were extracted. To characterize the spatial organization of FAs, cell masks were subdivided into ten concentric and contiguous rings, with ring 1 corresponding to the cell periphery and ring 10 corresponding to the nuclear region. Within each region, FAs were segmented using the FAAS and their distribution was calculated as the area covered by adhesions normalized to the total area of the corresponding ring.

### Cell tracking and migration analysis

Time-lapse sequences from both cell migration on uniform substrates and durotaxis on stiffness gradients were processed using a common custom MATLAB pipeline. Stage drift was corrected by tracking immobile debris within each field of view and subtracting their measured displacement from all frames. For durotaxis assays, the three overlapping DAPI frames spanning each stiffness gradient were first stitched in Fiji using the Grid/Collection Stitching plugin. The registration data generated by the plugin were used in MATLAB to stitch all frames from each position with identical *xy* coordinates. Nuclear centroids were detected with the TrackMate plugin (Fiji), and individual cell trajectories were reconstructed in MATLAB for quantification of migration parameters and angular trajectory distributions.

### Statistical analysis

All statistical analyses were conducted in R (v4.4.1) using custom scripts and standard libraries. The specific statistical tests applied are indicated in each figure. Non-parametric permutation and bootstrap methods were systematically used to ensure robust inference without distributional assumptions. Differences between groups or experimental conditions were assessed using permutation-based ANOVA models with 10,000 random resamplings. For single-factor analyses (e.g., effects of preconditioning on traction forces or migration parameters), one-way permutation ANOVA was applied. For multifactor analyses (e.g., effects of stiffness and preconditioning on traction forces or intracellular tension), two-way permutation ANOVA models were used, and *p*-values were computed for both linear and quadratic terms to test for biphasic stiffness dependencies. For directional migration analyses, two-sided one-sample permutation tests (10,000 resamples) were used to assess whether the mean *FMI_x_* differed significantly from zero, indicating directional bias. Paired permutation tests were used to compare migration directionality between conditions. Statistical significance was defined as *p* < 0.05. Data are represented as mean ± 95% confidence interval (bootstrap-estimated) or mean ± standard error of the mean (SEM), as specified in each figure.

### Cell accumulation

To determine whether the relative distribution of cells across stiffness regions changed between Day 0 and Day 2, we quantified the change in the fraction of cells within each stiffness interval for every gel (Δ*ρ* = *ρ*_*Day* 2_ − *ρ*_*Day* 0_). For each condition, the gels were divided into three stiffness regions (below, around, and above the peak). These boundaries were defined by identifying the position of the traction force peak and selecting adjacent stiffness ranges that maximized the slope of the traction force curve, thereby capturing the intervals of strongest durotaxis: below the peak (<1 kPa for plastic- and physiomimetic-preconditioned cells), around the peak (1–5 kPa for plastic- and 1–7 kPa for physiomimetic-preconditioned cells), and above the peak (≥5 kPa for plastic- and ≥7 kPa for physiomimetic cells). To analyze redistribution patterns while accounting for the non-independence of intervals measured within the same gel, we fitted linear mixed-effects models with stiffness interval as a fixed effect and gel identity as a random intercept. Planned contrasts were then used to directly compare the region around the peak with regions below and above it for each condition (e.g., Δ*ρ*_*Around the peak*_ − Δ*ρ*_*Below the peak*_, Δ*ρ*_*Around the peak*_ − Δ*ρ*_*Above the peak*_). Estimated marginal means and contrast statistics were obtained using the *emmeans* package, and two-tailed *p*-values were computed using Satterthwaite-approximated degrees of freedom.

## Acknowledgements

We thank all members of our groups for valuable discussions and continuous support. We are grateful to A. Faella for technical assistance and logistical support throughout the project, C. Herranz for support during the establishment of the laboratory and J. F. Abenza for providing reagents. We also thank P. Roca-Cusachs and X. Trepat for stimulating discussions. This work was funded in part by the Generalitat de Catalunya through AGAUR (2021 SGR 00523 to R.S., J.O. and R.F.) and by the Spanish Ministry of Science and Innovation (MICINN/FEDER; PID2024-159132OB-I00 and CNS2022-135533 to R.S., PID2021-128674OB-I00 to J.O. and R.S. and PID2023-146070OB-I00 to R.F.). Research reported in this publication was also supported by the National Institutes of Health (U54CA210190, P01CA254849, and U54CA268069) and by the National Science Foundation (grant no. 2222434). The content is solely the responsibility of the authors and does not necessarily represent the official views of the NIH or the NSF. M.M.L. and S.O. are supported by FPI predoctoral fellowships from the Spanish Agencia Estatal de Investigación (AEI; PRE2021-097143 and PRE2022-104818, respectively). R.S. is a Serra-Húnter Fellow.

## Declaration of Generative AI and AI-assisted technologies in the writing process

During the preparation of this work the authors used ChatGPT to improve the clarity of certain sentences. After using these tools, the authors reviewed and edited the content as needed and take full responsibility for the content of the publication. The use of AI tools was limited to improve writing and all ideas presented in this work remain original.

## Contributions

M.M.L. performed the experiments, analyzed the data, prepared the figures, and co-wrote the manuscript. R.A.M. performed the molecular clutch simulations and co-wrote the manuscript. S.O. contributed to the optimization of the experimental system and contributed to data interpretation. M.G.G. analyzed traction force microscopy data and computed intracellular stress fields. P.P. and R.F. contributed to data interpretation. J.O. developed the 3D lung-derived physiomimetic hydrogels. D.O. contributed to the simulations, data interpretation, and manuscript writing. R.S. conceived the project, designed the experiments and techniques, developed the phenomenological model, analyzed part of the data, and co-wrote the manuscript.

