## Supplementary Information / Extended Data for "Physiomimetic culture bias durotaxis toward soft environments"

### Supplementary Text

#### Supplementary Text I: Fabrication and measurement of polyacrylamide gels

Polyacrylamide (PAA) gels are the standard substrates for mechanobiology experiments, as they provide cells with mechanical environments of defined and tunable stiffness. PAA gels are obtained by polymerizing acrylamide and bis-acrylamide in water using ammonium persulfate (APS) as the initiator and tetramethylethylenediamine (TEMED) as the catalyst[1,2]. Stiffness is typically adjusted by changing the acrylamide-to-bis-acrylamide ratio. However, altering this ratio also modifies the polymer network structure with variable mesh sizes and porosity that may not scale with stiffness. This variability can produce changes in protein adsorption and cell behavior even at identical nominal stiffness. In addition, the literature reports a wide range of custom PAA formulations, often yielding different stiffness values even at similar acrylamide and bis-acrylamide concentrations, complicating cross-study comparisons.

To overcome these limitations, we adopted the formulation strategy introduced by Brock et al.[3], in which the crosslinker ratio ($C_m$) is held constant (0.02 mol mol$^{-1}$ bis-acrylamide to acrylamide) while the total monomer concentration is varied. This method generates a linear and predictable relationship between total monomer concentration and gel stiffness (Extended Data Fig. 1). Using the calibration reported by Brock et al., we extrapolated this relationship to obtain gels with target Young's moduli of 0.5, 1, 2.5, 5, 15, and 30 kPa (Extended Data Table 1). Gel stiffness was quantified by atomic force microscopy (AFM) indentation using spherical probes. Measurements obtained with pyramidal tips systematically overestimated the apparent modulus, likely due to nonlinear local deformations produced by sharp indenters[4]. In contrast, spherical tips distribute the applied load over a broader contact area, ensuring small strains and preserving linear elastic behavior. This configuration provides a more accurate estimate of the bulk modulus and enhances reproducibility across stiffness ranges. Taken together, fixing the crosslinker ratio while modulating stiffness through total monomer concentration, combined with AFM calibration using spherical tips, produced a reproducible and physically consistent PAA gels protocol for our study.

#### Supplementary Text II: Phenomenological simulation of cell migration on a stiffness gradient

To complement the experimental results shown in Fig. 3a-d and Extended Data Fig. 4, we implemented a phenomenological model of cell migration on a two-dimensional substrate with a stiffness gradient. This minimal framework was designed to reproduce the emergent redistribution of cells driven by durotaxis and to assess whether the experimentally observed values and trends are consistent with theoretical expectations. The simulations captured three qualitatively distinct regimes: positive durotaxis, corresponding to migration toward stiffer regions; negative durotaxis, corresponding to migration toward an optimal intermediate stiffness; and the absence of durotaxis, corresponding to random migration.

##### *Model overview*

Cells were modeled as point particles performing a persistent random walk with an optional durotactic bias. Each cell $i$ migrated within a rectangular domain representing the substrate, 2.5 mm in length and 1.5 mm in width (Extended Data Fig. 5a). The spatial profile of substrate stiffness, $E(x)$, varied along the $x$-axis according to a sigmoidal function fitted to AFM measurements of the stiffness profile of the PAA stiffness gradient (Extended Data Fig. 3b):

$$E(x) = \frac{25}{1 + \exp[-0.003(x - 1500)]}$$

where $E(x)$ is in kPa and $x$ (in µm) denotes the direction of increasing stiffness. This profile produces an effective gradient from 0 to 25 kPa across the gel.

For each cell $i$, the velocity evolves according to:

$$v_i(t) = \alpha\, v_i(t - \Delta t) + \sqrt{1 - \alpha^2}\, \eta_i + b_i(x)$$

where $\alpha$ is the persistence parameter, $\eta_i$ is a Gaussian random term, and $b_i(x)$ represents the durotactic bias.

##### *Durotactic bias*

The bias term $b_i(x)$ depends on the specific condition:

- **Positive durotaxis:** cells migrate preferentially toward stiffer regions. Multiple studies indicate that the magnitude of this directional bias diminishes as local stiffness increases[5,6]. To capture this behavior, we defined the stiffness-dependent bias using a sigmoid function that progressively reduces the bias strength with increasing stiffness (Extended Data Fig. 5b):

$$b_i(x) = L_{\text{low}} + \frac{b_0 - L_{\text{low}}}{1 + \exp\left[k_s \left(E(x) - x_{pos}\right)\right]} + \sigma_b \xi$$

  where $b_0$ is the mean bias, $L_{\text{low}}$ is the lower asymptotic value at high stiffness, $k_s$ controls the steepness of the transition, $x_{pos}$ is the stiffness midpoint of the sigmoid, $\sigma_b$the variability, and $\xi$ a Gaussian random variable.

- **Negative durotaxis:** cells display a biphasic durotactic response characterized by a smooth transition from positive durotaxis at low stiffness to negative durotaxis above an optimal stiffness. Cells migrate toward stiffer regions when $E(x) < 2.5$ kPa, exhibit negligible bias between approximately 2.5–5 kPa, and reverse their direction toward softer regions when $E(x) > 5$ kPa. This behavior is modeled with a continuous logistic function (Extended Data Fig. 5b):

$$b_i(x) = b_{\text{pos},0}\,[1 - s(a_1(E(x) - 1))] - b_{\text{neg},0}\, s(a_2(E(x) - x_{neg})) + \sigma_b\, \xi$$

where $b_{\text{pos},0}$ and $b_{\text{neg},0}$ are the mean biases for positive and negative durotaxis, respectively, $s(z) = \frac{1}{1+e^{-z}}$ is the sigmoid function, $a_1 = 2, a_2 = 10$ define the transition sharpness, $x_{neg}$ is the stiffness midpoint of the sigmoid, $\sigma_b$ is the standard deviation of the bias variability, and $\xi$ is a normally distributed random variable. This formulation captures the progressive shift from positive durotaxis toward the optimal stiffness and its subsequent reversal into negative durotaxis beyond that point, consistent with negative durotaxis experiments[7].

- **Random migration:** $b_i(x) = 0$, yielding an unbiased persistent random walk (Extended Data Fig. 5b).

In all cases, cells speed is reflected at the boundaries of the domain.

##### *Simulation procedure*

Each simulation represented 48 h of migration, discretized into 144 time steps of 20 min. For each substrate, 300 cells were simulated and initialized according to the experimentally measured distribution of *x*-positions at the initial time of the experiment. Twenty-five independent realizations were performed per durotactic condition. The stiffness gradient was partitioned into three regions (below the peak, around the peak and above the peak), and the fraction of cells within each region was quantified at time 0 and after 48 h. Changes in these fractions, expressed in percentage points (pp), quantified cell accumulation along the gradient. Parameters used in cell migration simulations can be found in Extended Data Table 3.

##### *Output metrics*

For each simulation, we computed:

- **Mean *x*-displacement ($\Delta x$):** *x*-displacement of the population centroid (i.e., the average of the individual net *x*-displacements of all cells) after 48 h, in µm.
- **Accumulation change ($\Delta\rho$):** variation in the percentage of cells below the peak, around the peak and above the peak regions in pp.

These simulations captured the qualitative features observed experimentally: accumulation toward stiff regions for positive durotaxis, accumulation toward optimal stiffness for negative durotaxis, and uniform distributions for random migration.

##### *Implementation details and reproducibility*

All simulations were performed in MATLAB R2023a (MathWorks, MA). Random number generation was initialized with a fixed seed (rng(25)) to ensure reproducibility of stochastic trajectories. Each condition (positive durotaxis, negative durotaxis, and random migration) was simulated independently using identical base code and parameter sets, differing only in the definition of the durotactic bias term. Reported results correspond to the mean of 25 independent realizations per condition.

*Results*

Simulations revealed distinct redistribution patterns that depended on the durotactic mode. Under negative durotaxis, cells progressively depleted both the soft and stiff regions and accumulated near the transition zone separating positive and negative durotaxis (Extended Data Fig. 5c). By contrast, positive durotaxis generated a pronounced depletion in the softer regions, followed by an enrichment at higher stiffness that diminished as the durotactic bias decreased (Extended Data Fig. 5d). In the absence of bias, random migration produced uniform cell distributions along the gradient (Extended Data Fig. 5e). Mapping spatial position $x$ into local stiffness values revealed that cells undergoing negative durotaxis accumulated near their optimal stiffness (Extended Data Fig. 5f), whereas positive durotaxis led to accumulation beyond this optimum (Extended Data Fig. 5g); no accumulation was observed for random migration (Extended Data Fig. 5h). In all cases, relative changes in cell density across regions remained below 5 pp.

To quantify directional bias independently of gel binning, we compared the population centroid at 0 and 48 h. The centroid shifted toward stiffer regions under positive durotaxis, toward softer regions under negative durotaxis, and remained unchanged in the absence of bias (Extended Data Fig. 5i). This centroid displacement therefore provides a robust measure of collective migration bias that is independent of gel binning.

#### Supplementary Text III: Physics-based Cell Migration Simulation

To interpret how culture preconditioning alters durotactic behavior across stiffness gradients, we developed a physics-based computational model that links molecular-scale force transmission to whole-cell migration. Experimental measurements revealed pronounced changes in cytoskeletal organization, adhesion architecture, traction forces, and durotactic directionality that could not be explained solely by static force-stiffness relationships. The goal of the model is therefore not to reproduce individual experimental observables in isolation, but to provide a unified mechanistic framework that connects actomyosin contractility, adhesion dynamics, and substrate mechanics to emergent migratory behavior. By explicitly resolving time-dependent force buildup, stochastic adhesion dynamics, and mechanical coupling across the cell, the model allows us to test how distinct motor-clutch regimes give rise to different durotactic responses under otherwise identical stiffness gradients. Detailed model formulation, assumptions, and simulation procedures are presented below.

##### *Model overview*

We model single-cell migration as an emergent mechanical process arising from stochastic force transmission between the actomyosin cytoskeleton and a deformable extracellular matrix. The model builds on the motor-clutch framework[8], in which myosin-driven actin retrograde flow is intermittently coupled to the substrate through transient adhesion bonds. Sensitivity to substrate stiffness is not imposed a priori, but arises naturally from the interplay between force generation, adhesion dynamics, and substrate deformation.

To connect molecular-scale force transmission to whole-cell migration, the cell is represented as a set of multiple peripheral force-generating units/modules elastically coupled to a central cell body (Fig. 4a). Each peripheral module generates active forces through actomyosin contraction and transmits traction forces to the substrate through adhesion clutches. In addition, the cell body contains a central force-transmitting unit that engages the substrate through adhesion clutches but does not generate active contractile forces, as no myosin motors are present in this unit. Instead, the central unit provides passive resistance to motion and enforces global force balance between peripheral traction forces and substrate coupling beneath the cell body. Together, elastic coupling between peripheral modules and the passive central unit ensures mechanical coherence and allows redistribution of internal forces across the cell.

Cell migration results from the balance of forces generated by these units as they stochastically engage, load, and disengage from the substrate in response to mechanical cues.

##### *Retrograde actin flow and motor-mediated force generation*

Within each force-generating unit, actin filaments undergo retrograde motion driven by myosin II motors. The velocity of actin flow decreases with increasing mechanical load and is described by a linear force-velocity relationship. Accordingly, the actin retrograde flow velocity of the $j$-th module is given by:

$$v_{act}^{j} = v_0\left(1 - \frac{F_c^{\,j}}{n_m^{j} F_m}\right) \tag{1}$$

where $v_0$ is the unloaded motor velocity, $F_c^{\,j}$ is the total force transmitted through engaged adhesions in the $j$-th force-generating unit, $n_m^j$ is the number of active motors in the $j$-th unit, and $F_m$ is the single-motor stall force. As force builds on engaged clutches, actin flow slows down and ultimately stalls when the motor ensemble reaches its load-bearing capacity (stall force).

#### *Actin polymerization and global actin conservation*

To regulate protrusion growth and retraction, actin turnover was explicitly incorporated into the model. F-actin polymerization within each module depends on the availability of monomeric actin, enforcing competition for a shared cellular pool. The instantaneous polymerization velocity is defined as:

$$v_p = v_p^* \frac{A_G}{A_T} \tag{2}$$

where $v_p^*$ denotes the maximal F-actin polymerization velocity, $A_G$ is the available G-actin in the cell, and $A_T$ is the total actin content. This formulation ensures global actin conservation and dynamically couples protrusion activity across modules.

#### *Adhesion clutches and force transmission*

Mechanical coupling between the actin cytoskeleton and the substrate is mediated by adhesion clutches that bind and unbind stochastically. Clutches bind with a force-independent on-rate $k_{on}$ and, when engaged, transmit force as linear springs, $F_c^{j,i} = \kappa_c x^{j,i}$, where $k_c$ is the clutch stiffness and $x^{j,i}$ is the extension of the $i$-th clutch in the $j$-th module. The total traction force generated by a module is given by the vector sum over all engaged clutches:

$$F_c^{\,j} = \sum_i F_c^{\,j,i} \tag{3}$$

The substrate beneath each unit is modeled as a linear elastic element with local effective stiffness $\kappa_{sub}$, such that:

$$F_c^{\,j} = \kappa_{sub} x_{sub}^j \tag{4}$$

where $x_{sub}^j$ is the substrate displacement in the $j$-th module. Each peripheral unit is elastically coupled to the cell body through a spring of stiffness $\kappa_{cell}$, transmitting a force of:

$$F_{cell}^j = \kappa_{cell} x_{cell}^j \tag{5}$$

where $x_{cell}^j$ and $\kappa_{cell}$ denote, respectively, the extension and constant of the Hookean spring connecting the cell body with the $j$-th module. In the central cell-body unit, clutches transmit force in the same manner, given by:

$$F_{cell} = \kappa_c \sum_{i=1}^{n_{c,cell}} x_{c,i} = \kappa_{sub} x_{sub,cell} \tag{6}$$

where $n_{c,cell}$ is the number of clutches associated with the central cell body, and $x_{sub,cell}$ is the substrate displacement in the cell body. In the cell body, actin retrograde flow is absent, and clutch loading arises solely from motion of the cell body relative to the substrate. The total cell traction force is computed as:

$$T = \sum_{j=1}^{N_{mod}} \left|F_c^{\,j}\right| + \left|F_{cell}^j\right| \tag{7}$$

Time-averaged traction forces are reported in dimensionless form by normalizing by the characteristic clutch module force $n_c^* F_b$ to allow better comparison between mechanical regimes.

Cell motion is governed by global force balance. According to Newton's second law, the net force exerted by the cell on the substrate satisfies, at all times:

$$\sum_{j=1}^{N_{mod}} F_c^{\,j} + F_{cell}^{\,j} = 0 \tag{8}$$

Directional motion arises from asymmetric force transmission across elastically coupled units, arising either from substrate stiffness gradients or from stochastic fluctuations in protrusion dynamics.

##### *Module nucleation and resource allocation*

Peripheral force-generating units were created stochastically at a rate that depends sensitively on the availability of actin monomers. The module nucleation rate is defined as:

$$k_{mod} = k_{mod}^{*} \left(\frac{A_G}{A_T}\right)^4 \tag{9}$$

where $k_{mod}^{*}$ is the maximal module nucleation rate under abundant G-actin conditions. This nonlinear dependence strongly suppresses protrusion formation when actin resources are limited. Upon formation, each module is initialized with fixed length $l^{mod}$ and assigned an orientation relative to the cell body. Motors and clutches are drawn from global pools of free motors $n_m^{free}$ and clutches $n_c^{free}$ according to:

$$n_m = n_m^{*} \frac{n_m^{free}}{n_m^{tot}}, \qquad n_c = n_c^{*} \frac{n_c^{free}}{n_c^{tot}} \tag{10}$$

where $n_m^{tot}$ and $n_c^{tot}$ are the total numbers of motors and clutches in the cell, and $n_m^{*}$ and $n_c^{*}$ are upper bounds on the number that can be recruited to a single module.

##### *Mechanosensitive motor recruitment*

In simulations incorporating mechanosensitive motors, the maximum number of motors that can be recruited to a newly nucleated module is not constant but depends on the local substrate stiffness sensed at the module position. Specifically, the upper bound on motor recruitment is defined as:

$$n_m^{*}(k_{sub}) = \alpha_m + \beta_m[2 + \log_{10}(k_{sub})] \tag{11}$$

where $\alpha_m$ sets the baseline motor recruitment level (upper bound on the number of motors for soft substrates, i.e. $k_{sub} = 0.01 \text{ pN nm}^{-1}$), and $\beta_m \geq 0$ quantifies the strength of motor mechanosensitivity. The non-mechanosensitive motor case corresponds to $\beta_m = 0$. This logarithmic dependence was motivated by the experimental observation that phosphorylated myosin levels increase with substrate stiffness (Fig. 1g-i). Motor recruitment therefore increases with substrate stiffness while remaining constrained by the availability of free motors in the global pool.

##### *Module capping and disassembly*

Modules are capped at rate $k_{cap}$, preventing further polymerization. Modules whose length falls below a minimum threshold $l^{min}$ are destroyed. Upon module destruction, all motors, clutches, and remaining actin are returned to their respective global pools.

##### *Force-dependent clutch dissociation and adhesion reinforcement*

Clutch dissociation is force-dependent and incorporates both slip-bond rupture and adhesion reinforcement. The dissociation rate of an individual clutch is given by:

$$k_{off}\left(\left|F_c^{j,i}\right|\right) = k_{off}^0 e^{\frac{\left|F_c^{j,i}\right|}{F_b\left(1+\alpha_{reinf}\left[\frac{n_c^j(F_c>F_{th})}{n_c^j}\right]^{\beta_{reinf}}\right)}} \tag{12}$$

where $k_{off}^0$ is the unloaded clutch off-rate, $F_b$ is its characteristic rupture force, $n_c^j$ is the total number of clutches in the $j$-th module, and $n_c^j(F_c > F_{th})$ is the number of clutches exceeding a threshold force $F_{th}$. The parameters $\alpha_{reinf}$ and $\beta_{reinf}$ control the strength and nonlinearity of reinforcement.

##### *Simulation procedure*

The stochastic dynamics are simulated using a direct Gillespie algorithm:

(1) Initialize a cell at a prescribed position with three identical modules of length $l^{mod}$, uniformly distributed in orientation, with no clutches bound, such that the system is initially force-free. In stiffness-gradients simulations, the initial position is set to the midpoint of the gradient.
(2) Compute rates for all possible events, including module nucleation, capping and disassembly, clutch binding and clutch unbinding.
(3) Determine the time to the next event based on the total event rate.
(4) Select and execute the event probabilistically.
(5) Update F-actin retrograde flow and actin polymerization velocities.
(6) Update module lengths, clutch extensions, and substrate displacements.
(7) Eliminate modules shorter than $l^{min}$.
(8) Enforce global force balance to determine cell body motion.
(9) Return to step 2.

Each durotaxis simulation was run for three hours of simulated time, with cell position, traction forces and module dynamics recorded throughout the simulation window.

##### *Substrate stiffness profile*

The substrate stiffness $k_{sub}(y)$ was defined as a one-dimensional, piecewise continuous function along the gradient axis $y$, consisting of two constant plateaus connected by smooth transition regions. The parameters $\kappa_{soft}$ and $\kappa_{stiff}$ denote the stiffness on the softer and stiffer sides of the gradient, respectively. The parameter $h_g$ denotes the width of the stiffness gradient.

- For $y \leq -h_g/2$, $\kappa_{sub}(y) = \kappa_{soft}$
- For $-h_g/2 < y \leq h_g/2$, $\kappa_{sub}(y) = \kappa_{soft} + 0.5\left[1 + erf\left(4y/h_g\right)\right]\Delta\kappa_{sub}$ (13)
- For $h_g/2 < y$, $\kappa_{sub}(y) = \kappa_{stiff}$

where $\Delta\kappa_{sub} = \kappa_{stiff} - \kappa_{soft}$. The error-function transition ensures smooth spatial variation of stiffness and avoids discontinuities in mechanical cues experienced by migrating cells.

##### *Consequences for stiffness-dependent force transmission*

Because clutch reinforcement depends on the accumulation of highly loaded clutches, its activation is inherently sensitive to substrate stiffness and force-loading rate. On compliant substrates, individual clutch forces remain small and clutch reinforcement is rarely engaged. On very stiff substrates, forces rise rapidly but clutch rupture occurs before sufficient reinforcement can develop. At intermediate stiffness, multiple clutches simultaneously exceed the force threshold, activating reinforcement and prolonging force transmission. This collective stabilization enhances traction generation near an optimal stiffness without altering the fundamental biphasic dependence of force on substrate stiffness.

#### Extended Data Figures

##### Extended Data Figure 1

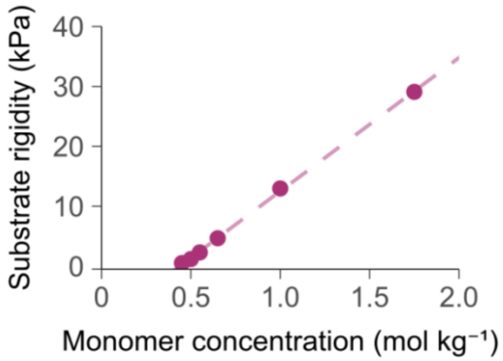


**Extended Data Figure 1. Calibration of PAA gel stiffness as a function of total monomer concentration**. Young's modulus of PAA gels as a function of total monomer concentration. Gels were prepared keeping the crosslinker ratio constant ($C_m$ = 0.02 mol mol$^{-1}$ bis-acrylamide to acrylamide). Each point represents the mean ± 95% confidence intervals estimated by bootstrap resampling of ≥80 AFM indentation measurements in 9 gels from three independent batches. The linear relationship obtained was used to interpolate acrylamide/bis-acrylamide concentrations corresponding to target stiffness values used in this study and shown in Extended Data Table 1.

Extended Data Figure 2

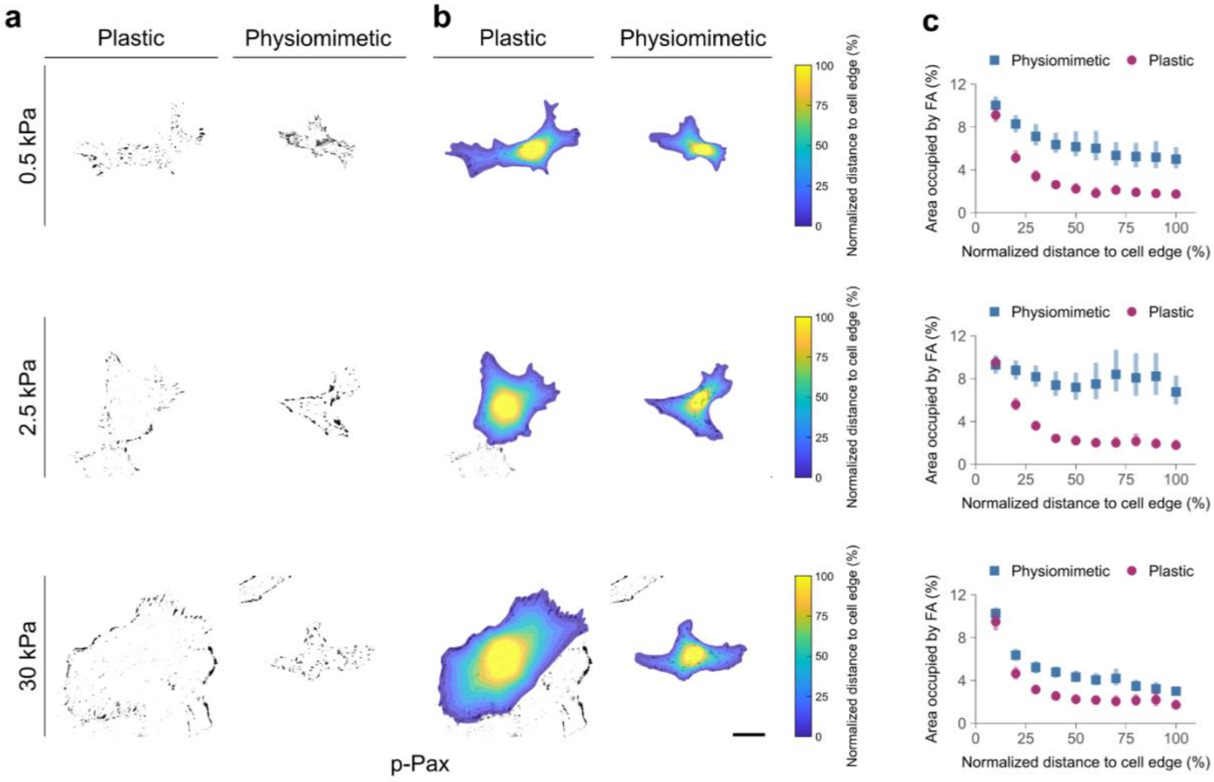


**Extended Data Figure 2. Focal adhesion (FA) localization. a,** Representative images of plastic- and physiomimetic-preconditioned cells seeded at 0.5, 2.5 and 30 kPa and stained for phosphorylated paxillin (p-Pax). Scale bar, 25 µm. **b,** Quantification of FA distribution. The cell spreading area was divided into ten concentric rings, and the percentage of area occupied by FAs was computed within each ring. **c,** Spatial distribution of FAs within cells. Plastic-preconditioned cells displayed a peripheral enrichment of FAs, whereas physiomimetic-preconditioned cells showed a more homogeneous distribution across the cell area at all rigidities. Data represent mean ± 95% confidence intervals estimated by bootstrap resampling of 45 cells from 3 independent experiments.

#### Extended Data Figure 3

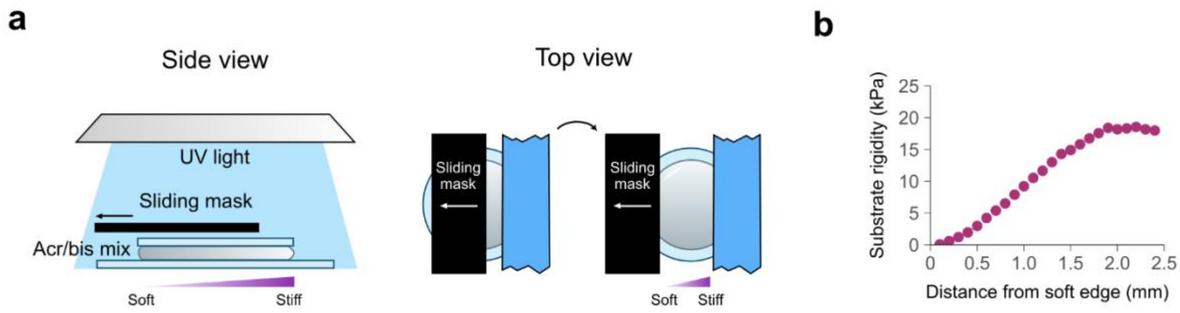


**Extended Data Figure 3. Fabrication of stiffness gradient gels. a,** To produce stiffness gradient gels, we irradiate an acrylamide/bis-acrylamide solution containing a photo initiator. Initially, the sample is protected by an opaque mask. An irradiation gradient is obtained by moving the mask (in the direction of the arrow) while illuminating the solution with a non-collimated UV lamp (365 nm). **b,** A 12% acrylamide and 1% bis-acrylamide solution produced a stiffness gradient gel of approximately 10 kPa $mm^{-1}$ when irradiated for 245 s moving the mask at 10 µm $s^{-1}$. Data represent mean ± 95% confidence intervals estimated by bootstrap resampling of 14 gels from three independent batches.

Extended Data Figure 4

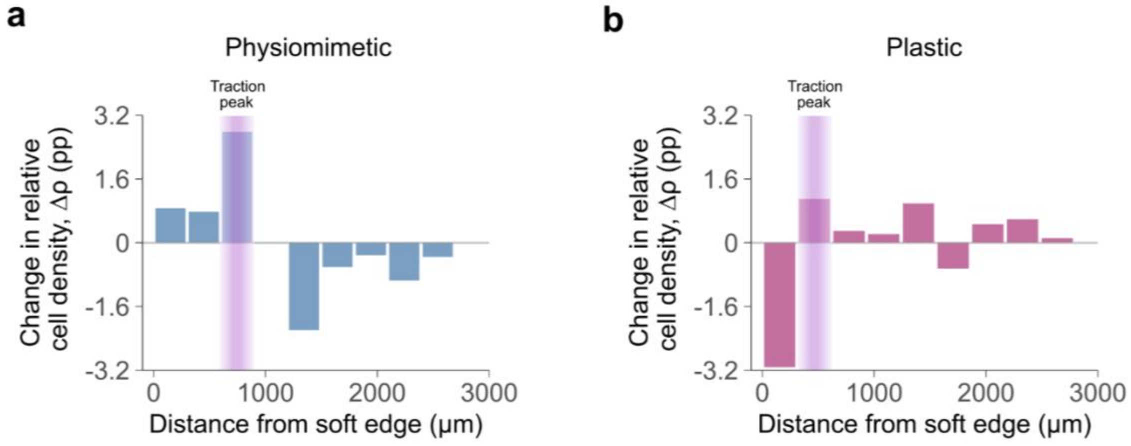


**Extended Data Figure 4. Change in cell density as a function of distance along the stiffness gradient.** Relative changes in local cell density ($\Delta\rho$), quantified in percentage points (pp), plotted as a function of distance along stiffness gradient gels spanning ~0.2–20 kPa. Positive $\Delta\rho$ values indicate cell accumulation, whereas negative $\Delta\rho$ values indicate depletion. Proliferation was arrested with thymidine, and phase-contrast images were acquired 4 h and 48 h after seeding (see Methods). **a,** Physiomimetic-preconditioned cells exhibited maximal accumulation near their optimal stiffness peak (≈600–900 µm) and a sharp decrease above the peak (>1200 µm), consistent with accumulation driven primarily by negative durotaxis from stiffer regions. **b,** In contrast, plastic-preconditioned cells depleted soft regions (<350 µm) and accumulated at higher stiffnesses, indicative of a positive-durotactic redistribution toward stiffer regions. Data are shown as $\Delta\rho$ relative to the initial distribution (n = 4,322 and n = 2,970 cells pooled from 7 independent gels for physiomimetic- and plastic-preconditioned cells, respectively).

#### Extended Data Figure 5

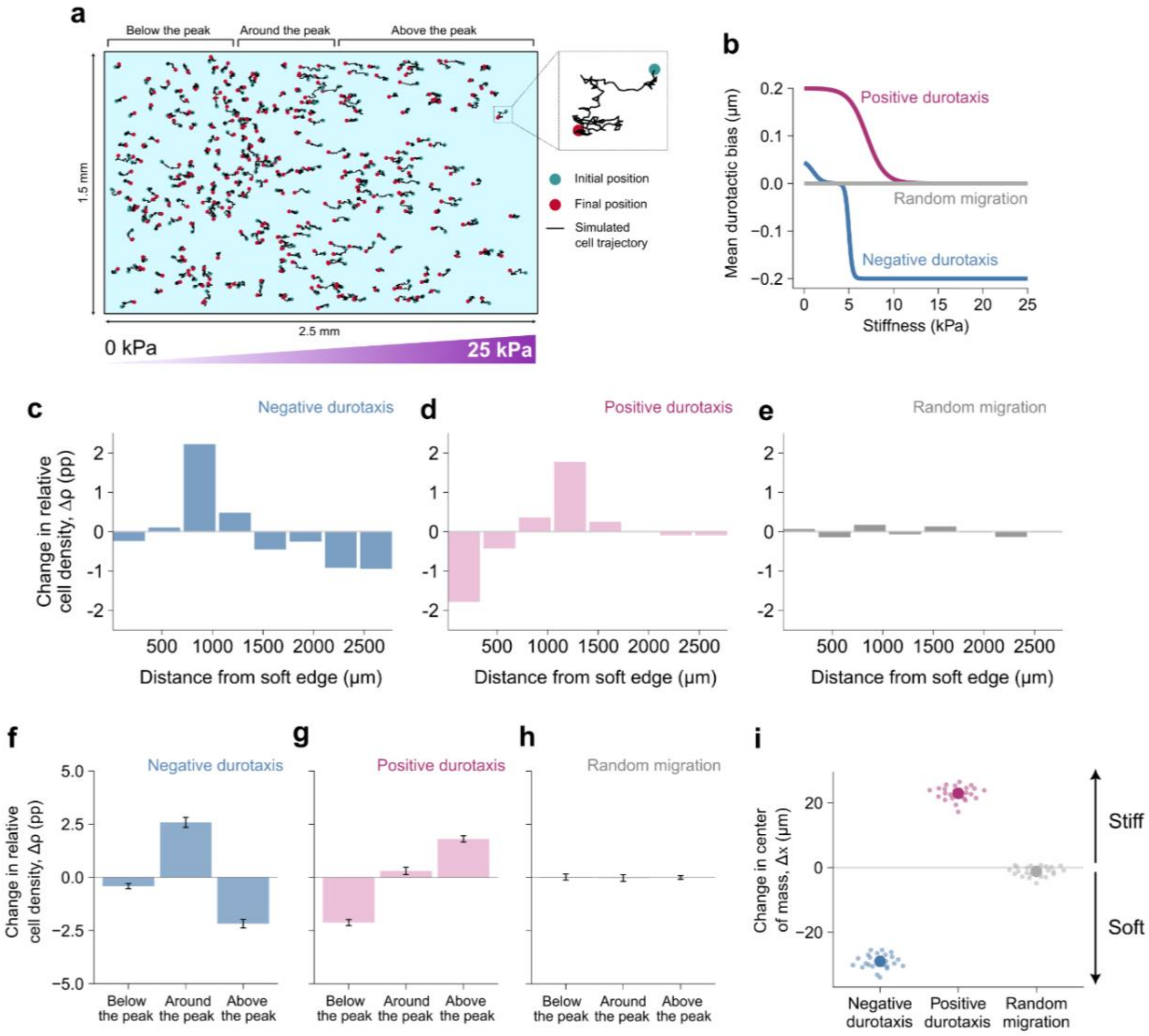


**Extended Data Figure 5. Simulated durotactic migration along a stiffness gradient. a,** Sketch of the phenomenological simulations. Cells are represented as active particles migrating on a two-dimensional substrate with a stiffness gradient ranging from 0 to 25 kPa. Each particle performs a persistent random walk with a tunable durotactic bias, enabling simulation of positive durotaxis (migration toward stiffer regions), negative durotaxis (migration toward an optimal stiffness), or unbiased migration (random migration). **b,** Mean durotactic bias as a function of stiffness. The mean durotactic bias varies with stiffness depending on the migration mode[5,9,10]. For positive durotaxis, the bias decreases as a sigmoid with stiffness (magenta line). For negative durotaxis, the bias is positive below, null around, and negative above the optimal stiffness $E^*$ (blue line). In the absence of durotaxis, the bias is set to zero (gray line). **c–e**, Spatial redistribution of cells as a function of distance along the stiffness gradient after 48 h of simulated migration. Bars indicate the change in the fraction of cells ($\Delta\rho$) in percentage points (pp). Negative durotaxis leads to accumulation at softer regions and a decrease in cell density at higher stiffnesses **(c)**, while positive durotaxis depletes softer regions **(d)**. Random migration shows no net redistribution **(e)**. **f–h**, Spatial redistribution of cells as a function of stiffness region after 48 h of simulated migration. Bars indicate the change in the fraction of cells ($\Delta\rho$ in pp) located below, around, or above the stiffness corresponding to the traction force peak. Negative durotaxis leads to accumulation near the optimal stiffness while a depletion below and above the peak **(f)**, positive durotaxis leads to cell accumulation around and beyond the stiffness peak **(g)**, and random migration shows no net redistribution **(h)**. Data represent mean ± SEM from 25 simulations. **i,** Comparison of cell positions at 0 and 48 h reveals that the population centroid shifts according to the durotaxis mode: toward stiffer regions under positive durotaxis, toward softer regions under negative durotaxis, and remains stationary in the absence of durotaxis.

#### Extended Data Figure 6

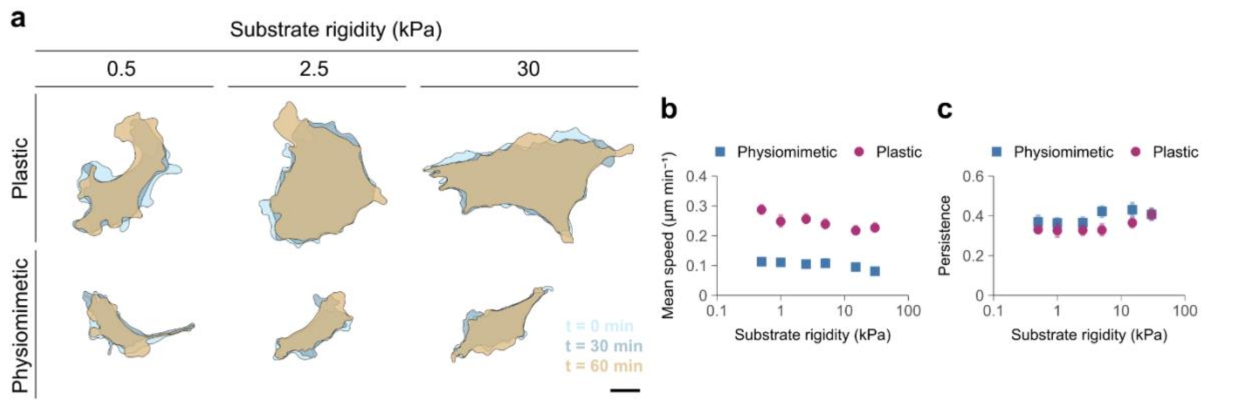


**Extended Data Figure 6. Migration dynamics on uniform stiffness substrates. a,** Dynamics (overlaid cell traces) of cells preconditioned on plastic or in physiomimetic hydrogels and subsequently seeded on uniform PAA gels of 0.5, 5 and 30 kPa. Overlaid cell contours are shown at 30-min intervals. Scale bar, 25 µm. **b,** Mean cell migration speed over >12 h on uniform stiffness gels (0.5–30 kPa). Images were acquired every 20 min. Speed decreased moderately with increasing rigidity. A permutation-based ANOVA (10,000 permutations) revealed significant effects of both substrate stiffness and preconditioning ($p < 0.0001$ for each), with plastic-preconditioned cells migrating on average ~2.5-fold faster than physiomimetic-preconditioned cells. **c,** Directional persistence of migration under the same conditions as in (**b**). Despite lower speeds, hydrogel-preconditioned cells exhibited significantly higher directional persistence than plastic-preconditioned cells (permutation-based ANOVA, $p < 0.0001$ for condition and stiffness effects). Data represent mean ± 95% confidence intervals estimated by bootstrap resampling of 85–251 cell trajectories from three independent experiments.

#### Extended Data Figure 7

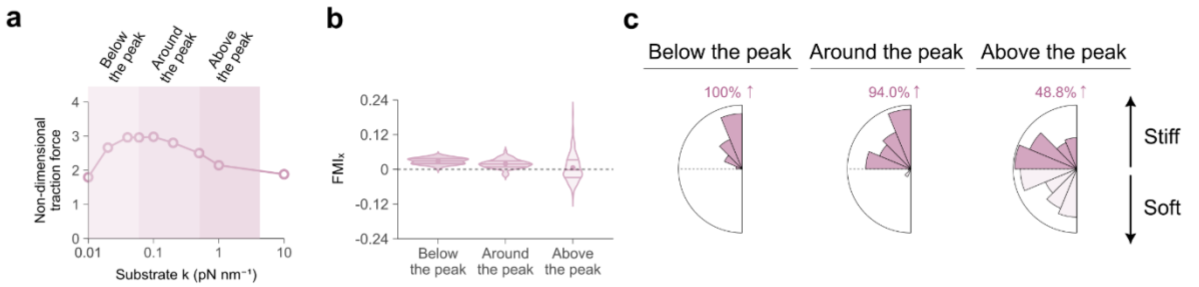


**Extended Data Figure 7. Molecular clutch model predicts that a high motor-high clutch regime suppresses negative durotaxis independently of mechanosensitivity. a,** Dimensionless total traction force as a function of substrate stiffness for the high motor-high clutch regime. A biphasic traction profile emerges, defining stiffness regions below the peak, around the peak, and above the peak. **b,** Forward migration index ($FMI_x$) for cells migrating within each stiffness region. Cells exhibit positive durotaxis below and around the traction force peak, while $FMI_x$ is near zero above the peak, indicating largely adurotactic migration. These results show that explicit mechanosensitivity of motors or clutches is not required to suppress negative durotaxis, and that high motor and clutch numbers are sufficient. **c,** Angular distributions of migration direction below the traction force peak, around the peak, and above the peak (n = 53, 134, 250 cells). Migration is biased toward increasing substrate stiffness below and around the peak, whereas migration above the peak lacks a preferred direction. Numbers in pink indicate the percentage of cells whose net migration direction had a positive projection along the stiffness gradient (positive durotaxis). Model parameter values are provided in Extended Data Table 4.

Extended Data Figure 8

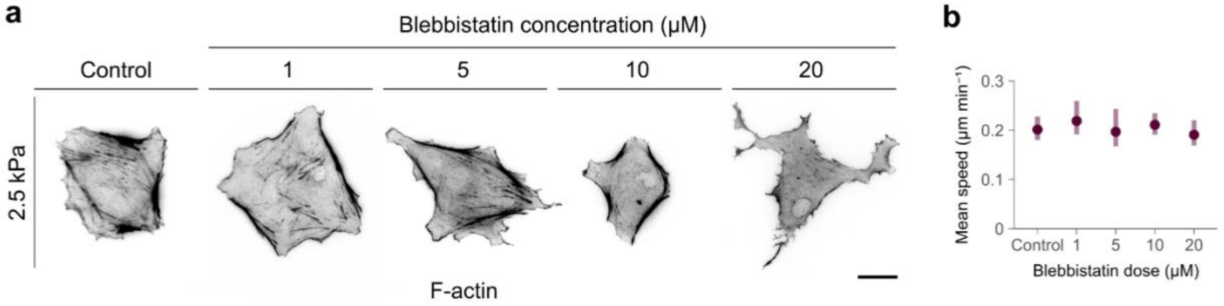


**Extended Data Figure 8. Effect of blebbistatin concentration on actin organization and cell migration.** **a,** Representative actin filaments (phalloidin) stainings of cells seeded on PAA gels of 2.5 kPa and treated with increasing concentrations of blebbistatin (1, 5, 10, and 20 µM). Higher doses progressively disrupted actin fiber organization. Scale bar, 25 µm. **b,** Average cell speed as a function of blebbistatin concentration. The 10 µM dose was selected for subsequent experiments as it induced clear cytoskeletal alterations without affecting migratory velocity. Data represent mean ± 95% confidence intervals estimated by bootstrap resampling of 39–102 cells from 1 independent experiment.

Extended Data Figure 9

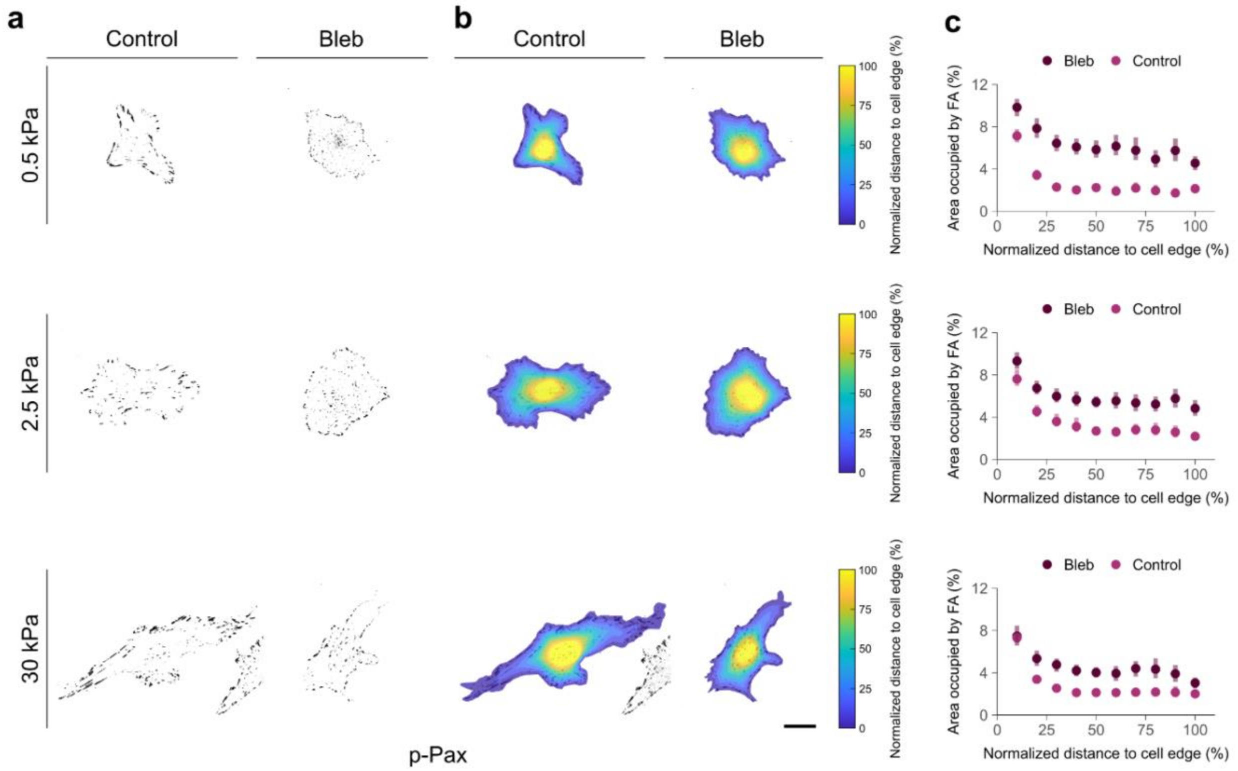


**Extended Data Figure 9. Focal adhesion (FA) organization in blebbistatin-treated cells. a,** Representative images of plastic-preconditioned cells seeded at 0.5, 2.5 and 30 kPa, treated with DMSO (Control) or 10 µM blebbistatin (Bleb) and stained for phosphorylated paxillin (p-Pax). Scale bar, 25 µm. **b,** Quantification of FA spatial distribution, performed as described in Extended Data Fig. 2. **c,** Spatial distribution of FAs within cells. Control cells displayed a clear peripheral enrichment of FAs, whereas blebbistatin-treated cells showed a more uniform distribution across the cell area. Data represent mean ± 95% confidence intervals estimated by bootstrap resampling of 30–57 cells from 3 independent experiments.

### Extended Data Tables

#### Extended Data Table 1

| Monomer concentration (mol kg$^{-1}$) | Acrylamide 40% (µl) | Bis-acrylamide 2% (µl) | $H_2O$ (µl) | Nominal stiffness (kPa) | Experimental stiffness (kPa) |
|---|---|---|---|---|---|
| 0.45 | 77.9 | 68.9 | 847.7 | 0.5 | 0.53 ± 0.1 |
| 0.5 | 81.7 | 72.3 | 840.5 | 1 | 1.20 ± 0.15 |
| 0.55 | 93.1 | 82.4 | 819 | 2.5 | 2.30 ± 0.45 |
| 0.65 | 112 | 99.1 | 783.4 | 5 | 4.60 ± 0.05 |
| 1 | 187.8 | 166.3 | 640.4 | 15 | 13.97 ± 1.47 |
| 1.75 | 301.4 | 266.9 | 426.2 | 30 | 29.09 ± 2.14 |

**Extended Data Table 1. PAA gel formulations corresponding to target stiffness values.** The linear calibration curve of Extended Data Fig. 1 was used to obtain gels with target Young's modulus of 0.5, 1, 2.5, 5, 15, and 30 kPa. Stiffness values were validated by AFM indentation with spherical probes to ensure consistency and reproducibility across preparations. Data are mean ± SD.

#### Extended Data Table 2

| Acrylamide 40% (μl) | Bis-acrylamide 2% (μl) | $H_2O$ (μl) | 10× PBS (μl) | Irgacure 2959 (μl) |
|---|---|---|---|---|
| 295.5 | 133 | 458 | 98.5 | 15 |

**Extended Data Table 2. PAA gel formulation for stiffness gradient gels.** The system of Extended Data Fig. 3 was used to obtain gels with a stiffness gradient ranging from ~0.2 to 20 kPa. Stiffness values were validated by AFM indentation with spherical probes to ensure consistency and reproducibility across preparations.

#### Extended Data Table 3

| Parameter | Symbol | Value | Pos./neg./rand. | Description |
|---|---|---|---|---|
| **Domain length** | $L$ | 2500 µm | All simulations | Length of the stiffness gradient |
| **Domain width** | $W$ | 1500 µm | All simulations | Width of the stiffness gradient |
| **Number of cells** | $N$ | 300 | All simulations | Number of cells per simulation |
| **Simulation duration** | $T$ | 48 h | All simulations | Total simulated time |
| **Time step** | $\Delta t$ | 20 min | All simulations | Integration interval |
| **Cell speed** | $v$ | 0.15 µm $min^{-1}$ | All simulations | Average migratory speed of each cell |
| **Persistence parameter** | $\alpha$ | 0.3 | All simulations | Controls the degree to which cells retain their previous migration direction |
| **Bias variability** | $\sigma_\beta$ | 0.8 | Pos./Neg. Sim. | Standard deviation of durotactic bias |
| **Optimal stiffness** | $E^*$ | 5 kPa | Neg. Simulations | Target stiffness for neg. durotaxis |
| **Mean bias strength** | $b_0$ | 0.2 µm | Pos. Simulations | Mean magnitude of stiff-directed bias |
| **Lower asymptotic bias value** | $L_{low}$ | 0 | Pos. Simulations | Minimal bias at high stiffness in the sigmoid formulation |
| **Sigmoid steepness** | $k_s$ | 1 | Pos. Simulations | Steepness of the stiffness-dependent transition in the bias |
| **Positive durotaxis midpoint** | $x_0$ | 7 kPa | Pos. Simulations | Stiffness at which the positive durotaxis decreases sharply |
| **Positive durotaxis bias** | $b_{pos,0}$ | 0.05 µm | Neg. Simulations | Mean durotactic bias below $E^*$ |
| **Negative durotaxis bias** | $b_{neg,0}$ | 0.2 µm | Neg. Simulations | Mean durotactic bias above $E^*$ |
| **Sigmoid sharpness parameter 1** | $a_1$ | 2 | Neg. Simulations | Controls the soft-side transition in the logistic bias function |
| **Sigmoid sharpness parameter 2** | $a_2$ | 10 | Neg. Simulations | Controls the soft-side transition in the logistic bias function |
| **Neg. durotaxis bias** | $x_{neg}$ | 5 kPa | Neg. Simulations | Stiffness at which the negative durotaxis increases sharply |
| **Gradient stiffness range** | – | 0-25 kPa | Pos./Neg. Sim. | Experimental stiffness profile |

**Extended Data Table 3. Parameters used in the phenomenological cell migration simulations.**

#### Extended Data Table 4

| Parameter | Symbol | Value (Cell 1*) | Value (Cell 2**) | Value (Cell 3***) |
|---|---|---|---|---|
| **Total number of motors** | $n_m^{tot}$ | $10n_m^*$ | 1000 | 10000 |
| **Single motor stall force** | $F_m$ | 2 pN | 2 pN | 2 pN |
| **Baseline motor recruitment level** | $\alpha_m$ | 1000 | 100 | 1000 |
| **Strength of motor mechanosensitivity** | $\beta_m$ | 300 | 0 | 0 |
| **Unloaded motor velocity** | $v_0$ | 120 nm $s^{-1}$ | 120 nm $s^{-1}$ | 120 nm $s^{-1}$ |
| **Total number of clutches** | $n_c^{tot}$ | 7500 | 750 | 7500 |
| **Maximum module number of clutches** | $n_c^*$ | 75 | 75 | 750 |
| **Clutch binding rate** | $k_{on}$ | 0.3 $s^{-1}$ | 0.3 $s^{-1}$ | 1 $s^{-1}$ |
| **Unloaded clutch dissociation rate** | $k_{off}^0$ | 0.1 $s^{-1}$ | 0.1 $s^{-1}$ | 0.1 $s^{-1}$ |
| **Strength of clutch reinforcement** | $\alpha_{reinf}$ | 1 | 0 | 0 |
| **Nonlinear exponent of clutch reinforcement** | $\beta_{reinf}$ | 0.5 | 0 | 0 |
| **Threshold force for clutch reinforcement** | $F_{th}$ | 2 pN | $\infty$ | $\infty$ |
| **Clutch spring constant** | $\kappa_c$ | 0.8 pN $nm^{-1}$ | 0.8 pN $nm^{-1}$ | 0.8 pN $nm^{-1}$ |
| **Characteristic clutch rupture force** | $F_b$ | 2 pN | 2 pN | 2 pN |
| **Substrate spring constant** | $\kappa_{sub}$ | Variable | Variable | Variable |
| **Substrate stiffness gradient width** | $h_g$ | $\frac{\Delta\kappa_{sub}}{3\cdot10^5}$ [nm] | $\frac{\Delta\kappa_{sub}}{3\cdot10^5}$ [nm] | $\frac{\Delta\kappa_{sub}}{3\cdot10^5}$ [nm] |
| **Total amount of actin in the cell** | $A_T$ | $10^5$ nm | $10^5$ nm | $10^5$ nm |
| **Maximum F-actin polymerization velocity** | $v_p^*$ | 200 nm $s^{-1}$ | 200 nm $s^{-1}$ | 200 nm $s^{-1}$ |
| **Maximum module birth rate** | $k_{mod}^*$ | 1 $s^{-1}$ | 1 $s^{-1}$ | 1 $s^{-1}$ |
| **Module capping rate** | $k_{cap}$ | $10^{-3}$ $s^{-1}$ | $10^{-3}$ $s^{-1}$ | $10^{-3}$ $s^{-1}$ |
| **Initial module length** | $l^{mod}$ | 100 nm | 100 nm | 100 nm |
| **Cell spring constant** | $\kappa_{cell}$ | $10^3$ pN $nm^{-1}$ | $10^3$ pN $nm^{-1}$ | $10^3$ pN $nm^{-1}$ |
| **Number of cell body clutches** | $n_{c,cell}$ | 100 | 10 | 100 |

Cell 1*: high motor cell with mechanosensitive motors and adhesion reinforcement

Cell 2**: low motor, low clutch cell

Cell 3***: non-mechanosensitive, high motor, high clutch cell

**Extended Data Table 4. Model parameters used in the cell migration motor-clutch simulations.**

#### Supplementary Videos legends

**Supplementary Video 1**. Phase-contrast time-lapse imaging of RFL-6 fibroblasts preconditioned on tissue-culture plastic or within physiomimetic 3D hydrogels and subsequently plated on PAA gels of uniform stiffness (0.5, 5, 30 kPa). Images were acquired for 1 h at 1 s intervals. Plastic-preconditioned cells display extensive lamellipodial activity and larger displacements, whereas hydrogel-preconditioned cells exhibit reduced protrusive dynamics and more persistent trajectories. Scale bar, 25 µm.

**Supplementary Video 2**. Phase-contrast time-lapse imaging (20 min per frame, 12 h total) of fibroblasts preconditioned on plastic or in physiomimetic hydrogels migrating on PAA gels of 0.5, 2.5 and 30 kPa. Plastic-preconditioned cells migrate faster and follow more variable trajectories, whereas physiomimetic-preconditioned cells migrate slower but with greater directional persistence. Scale bar, 100 µm.
